# Restricted MHC-II trafficking in *Mycobacterium tuberculosis*-infected M2-like macrophages limits CD4+ T cell activation

**DOI:** 10.64898/2026.07.07.736943

**Authors:** Avinaash K. Sandhu, Daniel P. Gail, Rachel C. Simmermon, Demetria Webb, Laura Hmiel, Charles Bark, Bryan Bryson, Richard F. Silver, Stephen Carpenter

## Abstract

Recognition of infected macrophages by CD4+ T cells is essential to immune protection against *Mycobacterium tuberculosis* (Mtb), the causative agent of tuberculosis (TB). However, not all infected macrophage subsets successfully elicit T cell activation. We recently discovered that M2-like macrophages fail to efficiently activate memory CD4+ T cells when infected with Mtb, yet successfully elicit T cell activation when loaded with peptides, γ-irradiated bacteria, or Mtb whole cell lysate. Since the mechanisms underlying CD4+ T cell evasion by infected M2 but not M1-like macrophages remain underexplored, we sought to determine the genes and pathways unique to Mtb infection of M2-like cells, including alveolar macrophages. RNA sequencing of human macrophages infected with virulent Mtb identified enrichment of IL-10 and type I interferon (IFN) signaling genes, including *IL10RA* and *HERC5*, respectively, in infected M2-like monocyte-derived and alveolar macrophages. However, genes involved in MHC-II trafficking, such as *AP1M2*, were higher in infected M1-like macrophages. In complementary experiments using fluorescence microscopy and flow cytometry, we observed impaired trafficking of newly synthesized MHC-II to the plasma membrane of Mtb-infected M2-like macrophages despite high total surface MHC-II levels. Neutralization of IL-10 or knockdown of *HERC5* restored MHC-II trafficking to the cell surface among infected M2-like macrophages and significantly enhanced activation of memory CD4+ T cells in an MHC-II-dependent manner. These findings identify coordinated IL-10 and type I IFN signaling as key mechanisms that restrict MHC-II trafficking to the plasma membrane in Mtb-infected M2-like macrophages, thereby limiting antigen presentation and CD4+ T cell activation. We propose that host-directed therapies targeting these pathways in infected alveolar macrophages will facilitate T cell recognition for the prevention or treatment of active TB.

**Author Summary:** Recognition of infected macrophages by CD4+ T cells is essential to immune protection against *Mycobacterium tuberculosis* (Mtb), the causative agent of tuberculosis (TB). However, not all infected macrophage subsets successfully elicit T cell activation. We recently discovered that M2-like macrophages fail to efficiently activate memory CD4^+^ T cells when infected with Mtb, yet they successfully elicit T cell activation when treated with peptides, γ-irradiated bacteria, or Mtb whole cell lysate. In this study, we identified genes and pathways uniquely upregulated in Mtb-infected M2-like macrophages that are linked to inefficient CD4+ T cell activation, including IL-10 signaling and type I interferon (IFN) pathways. These pathways were linked to reduced MHC-II trafficking to the plasma membrane in Mtb-infected M2-like macrophages. Neutralization of IL-10 or knockdown of *HERC5* restored MHC-II trafficking and augmented memory CD4+ T cell activation. Our study demonstrates that IL-10 signaling and type I IFN pathways play detrimental roles in macrophages during Mtb infection, impairing MHC-II trafficking and CD4+ T cell activation. Since lung-resident alveolar macrophages express a dominant M2-like phenotype, these findings suggest that targeting IL-10 and type I IFN signaling may offer a strategy to enhance CD4+ T cell-mediated immunity and improve TB outcomes.

## Introduction

Tuberculosis (TB) remains a leading global health threat, with over 10.8 million incident cases and over 1.2 million deaths in 2024 [1]. CD4+ T cells are critical for protection against TB [2]. Protection is conferred by T cell antigen receptor (TCR) recognition of peptide-bound major histocompatibility complex (pMHC) on the surface of infected macrophages [3–5]. Successful T cell recognition depends on the processing of *Mycobacterium tuberculosis* (Mtb) proteins and presentation of peptides in the context of MHC molecules, dynamics that vary across macrophage subsets [6,7]. Pro-inflammatory (M1-like) macrophages more efficiently restrict bacterial growth, while alternatively activated/anti-inflammatory (M2-like) macrophages are more frequently infected and are permissive to Mtb growth [6,8]. We recently showed that M2-like macrophages fail to efficiently activate human CD4+ T cells upon Mtb infection, while M1- and M2-like macrophage subsets both activated T cells when treated with exogenous antigens in the form of peptides, γ-irradiated bacteria, or Mtb whole cell lysate [9]. Together, these findings suggest that Mtb thrives within M2-like macrophages, evading T cell responses. However, the mechanisms by which Mtb leverages the environment of these antigen-presenting cells (APCs) to prevent T cell recognition remain unknown.

Several lines of evidence indicate that T cells inefficiently recognize Mtb-infected lung-resident alveolar macrophages (AMs). In the mouse model of TB, AMs are the only cells found to harbor bacilli for nearly two weeks after aerosol Mtb infection, at which point monocyte-derived macrophages (MDMs), dendritic cells (DCs), and neutrophils are recruited and become infected [10–13]. T cell responses in the lungs are similarly delayed by up to two weeks after aerosol infection [14,15]. Memory T cell responses are also delayed [4,16,17]. In humans, conversion of the tuberculin skin test or interferon-gamma release assay (IGRA), indicating Mtb-specific T cell responses, is delayed by up to 8-12 weeks after exposure to a household contact with TB [18,19]. Differences between murine and human macrophage responses to Mtb infection may account for certain gaps in our understanding of immunity to TB. In mice, Mtb infection reduces MHC-II expression through downregulation of the MHC class II transactivator (CIITA), limiting T cell responses [20]. In contrast, human M2-like MDMs show greater cell surface expression of the MHC-II gene HLA-DR compared to M1-like cells after Mtb infection, despite failing to efficiently activate CD4+ T cells [9,21]. Human AMs obtained from bronchoalveolar lavage (BAL) samples share phenotypic features with M2-like human MDMs, including the ability to respond to IL-10 [8,9,22–25]. Why human M2-like macrophages fail to activate CD4+ T cells during Mtb infection despite high surface MHC-II expression, and the role of IL-10, remain poorly understood.

IL-10 is an anti-inflammatory cytokine abundant in healthy lung tissue, while type I interferons (IFN) are induced by pattern recognition receptor ligation, typically during viral infection [9,26,27]. Yet both are considered detrimental to protection against early Mtb infection [10,13,28–34]. IL-10 was shown to impair phagosome maturation in Mtb-infected human MDMs and AMs, resulting in reduced bacterial killing and increased burden [33], and to reduce Th1 cell recruitment into the lung [34]. Furthermore, neutralization of IL-10 in CBA/J mice was associated with greater control of Mtb infection, increased T cell infiltration into the lungs, and enhanced IFN-ү production by T cells [28]. IL-10 has been shown to impair antigen presentation in professional APCs in multiple ways, including by reducing MHC-II surface expression and through induction of MARCH1, which ubiquitinates HLA-DR, targeting it for degradation [35]. Prior work demonstrated that IL-10 pre-treatment of human MDMs before Mtb infection resulted in retention of pMHC complexes in endosomal compartments upon Mtb infection, reducing MHC-II trafficking to the plasma membrane [36]. Interestingly, type I IFNs can drive increased IL-10 production by APCs and T cells [37–39]. Importantly, type I IFN responses are elevated in patients with active TB, compared to those with latent Mtb infection (LTBI) [29,40,41]. Knockout of the type I IFN receptor (IFNAR) rescued the neutrophil-driven inflammatory response to Mtb in Sp140^-/-^ mice, and improved IFN-ү receptor expression, highlighting the detrimental effects of type I IFN on control of Mtb infection [13]. Because IFN-γ signaling is intimately tied to increased MHC-II expression via induction of CIITA, type I IFNs can directly reduce MHC-II expression by APCs through decreasing surface *IFNGR* expression and IFN-γ signaling [42]. Taken together, IL-10 signaling, alone or in concert with type I IFN, can reduce antigen presentation in Mtb-infected macrophages.

In this study, we aimed to identify the key pathways responsible for subversion of the CD4+ T cell response by Mtb-infected M2-like macrophages. Using bulk-RNA sequencing to compare Mtb-infected M1 and M2-like MDMs, we found that Mtb-infected M2-like macrophages upregulated key genes in the type I IFN and IL-10 signaling pathways, including *HERC5* and *IL10RA,* respectively. These genes contributed to impaired MHC-II trafficking to the plasma membrane of Mtb-infected M2-like macrophages. Knockdown of *HERC5*, a key type I IFN stimulated gene, and neutralization of IL-10 in infected M2-like macrophages each resulted in increased trafficking of MHC-II to the cell surface. Critically, *HERC5* knockdown alone and in combination with IL-10 blockade enhanced recognition of Mtb-infected M2-like macrophages by Mtb-specific CD4+ T cells. This study highlights the roles of type I IFN and IL-10 signaling in impaired antigen presentation to CD4+ T cells through reduced MHC-II trafficking.

## Results

### Alveolar macrophages infected with Mtb do not efficiently activate CD4+ T cells

Lung-resident alveolar macrophages represent a cellular niche for Mtb infection in humans [6,12]. We and others find AMs to represent >85% of CD45+ adherent cells from human BAL samples, and express a CD169^Hi^ CD11c^Hi^ CD11b^Lo/Int^ CD163^Hi^ CD86^Hi^ phenotype similar to M2-like MDMs, while also displaying CD206^Hi^ and CD14^Lo^ surface marker expression akin to M1-like macrophages [9,22]. Furthermore, single-cell transcriptomics data for healthy human lung in the Genotype-Tissue Expression (GTEx) portal indicates both *CSF1* (M-CSF) and *CSF2* (GM-CSF) are expressed by alveolar epithelial cells, suggesting that AMs are exposed to both cytokines [9]. To examine the extent to which Mtb-infected human AMs activate autologous CD4+ T cells, we obtained BAL and peripheral blood samples from healthy individuals with LTBI and compared T cell activation in response to Mtb-infected BAL macrophages and MDMs. CD14+ monocytes were isolated by positive immunomagnetic selection from PBMCs and differentiated using GM-CSF or M-CSF, as described [9,43]. BAL cells were plated on the final day of MDM differentiation (Day 6) and both were infected with Mtb (strain H37Rv) at a multiplicity of infection (MOI) of 4 which was previously shown to infect >70% of M1 and >90% of M2-like MDMs [9] **(Fig. 1A)**. Following a 4 h infection and washout of extracellular bacteria, non-adherent BAL cells (containing T cells), or peripheral blood memory (CD45RA^Lo^) CD4+ T cells isolated by negative immunomagnetic selection, were co-cultured with macrophages infected with Mtb, treated with H37Rv whole cell lysate (Mtb lysate), or controls conditions. After 16-18 h, T cell recognition was quantified using activation-induced markers (AIMs) as previously reported [9,43,44] **(Fig. 1A)**. Both peripheral blood and BAL memory CD4^+^ T cells co-expressed CD69 and CD40L in response to Mtb-infected M1-like MDMs, compared to α-MHC-II mAb blockade **(Figs. 1B, 1C)**. However, significantly fewer peripheral blood or BAL CD4+ T cells were activated in response to BAL macrophages **(Figs. 1B-E).** Yet memory CD4+ T cells were successfully activated in response to M2-like MDMs pre-treated with Mtb lysate **(Fig. 1B)**, consistent with our prior results [9]. Therefore, similar to M2-like MDMs, memory CD4+ T cells do not efficiently recognize human AMs upon Mtb infection.

**Figure 1.**
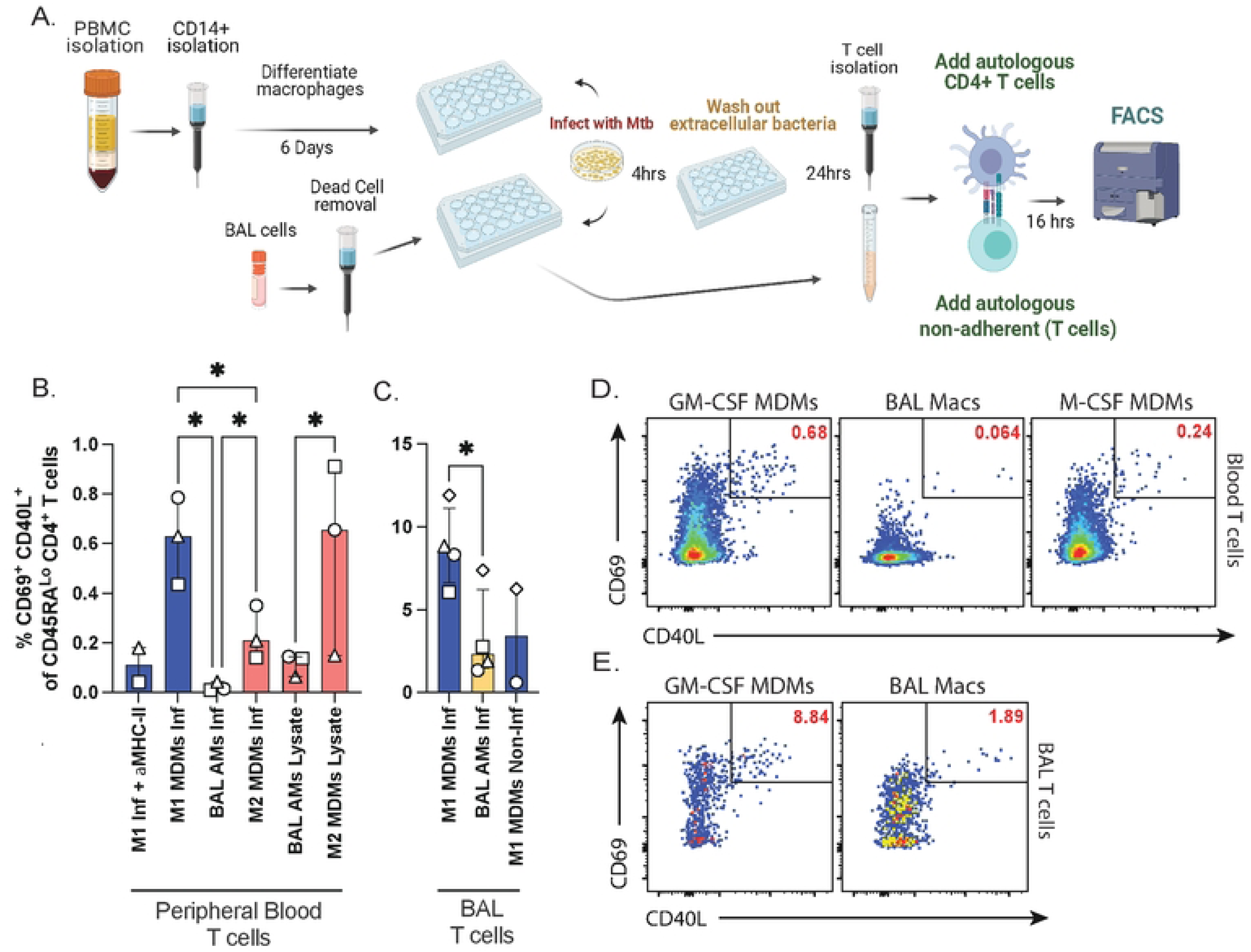
Alveolar macrophages infected with Mtb do not efficiently activate CD4+ T cells. **(A)** Schematic of experimental workflow to coculture infected M1-like or BAL macrophages with autologous memory CD4+ T cells or BAL T cells for flow cytometry. Created in BioRender. **(B)** Summary bar graphs comparing median (and inter-quartile range) co-expression of CD69 and CD40L gated on CD45RA^Lo^ (memory) peripheral blood CD4+ T cells or **(C)** BAL T cells after 16-18 h in co-culture with Mtb-infected MDMs and BAL macrophages. **(D)** Representative flow cytometry plots of CD69 and CD40L co-expression for memory CD4+ T cells from peripheral blood, or **(E)** BAL, 16-18 h after co-culture with Mtb-infected M1 or M2-like MDMs, or BAL macrophages. Data represent 1-2 replicates from 3-5 individual donors (independent experiments). Significance was determined using a paired mixed-effects one-way ANOVA with Geisser-Greenhouse correction and Šídák’s post test, corrected for multiple comparisons. ∗ p < 0.05; ns, not significant.

### Alveolar macrophages show enrichment of key type I IFN signaling genes and *IL10RA* upon Mtb-infection

Given the reduced CD4+ T cell responses to infected M2-like MDMs and AMs, we next sought to determine the genes and pathways shared by and unique to both macrophage subsets upon Mtb infection. Bulk RNA sequencing was performed on MDM and BAL macrophage lysates 24 h after Mtb infection or incubation with media. Genes upregulated by Mtb-infected vs. non-infected M1 or M2-like MDMs were compared to those upregulated by infected vs. non-infected BAL macrophages **(Fig. 2A)**. A small subset of 35 differentially-expressed genes (DEGs) was common only to infected M2-like MDMs and BAL macrophages, while 293 DEGs were upregulated by all 3 infected macrophage subsets.

**Figure 2.**
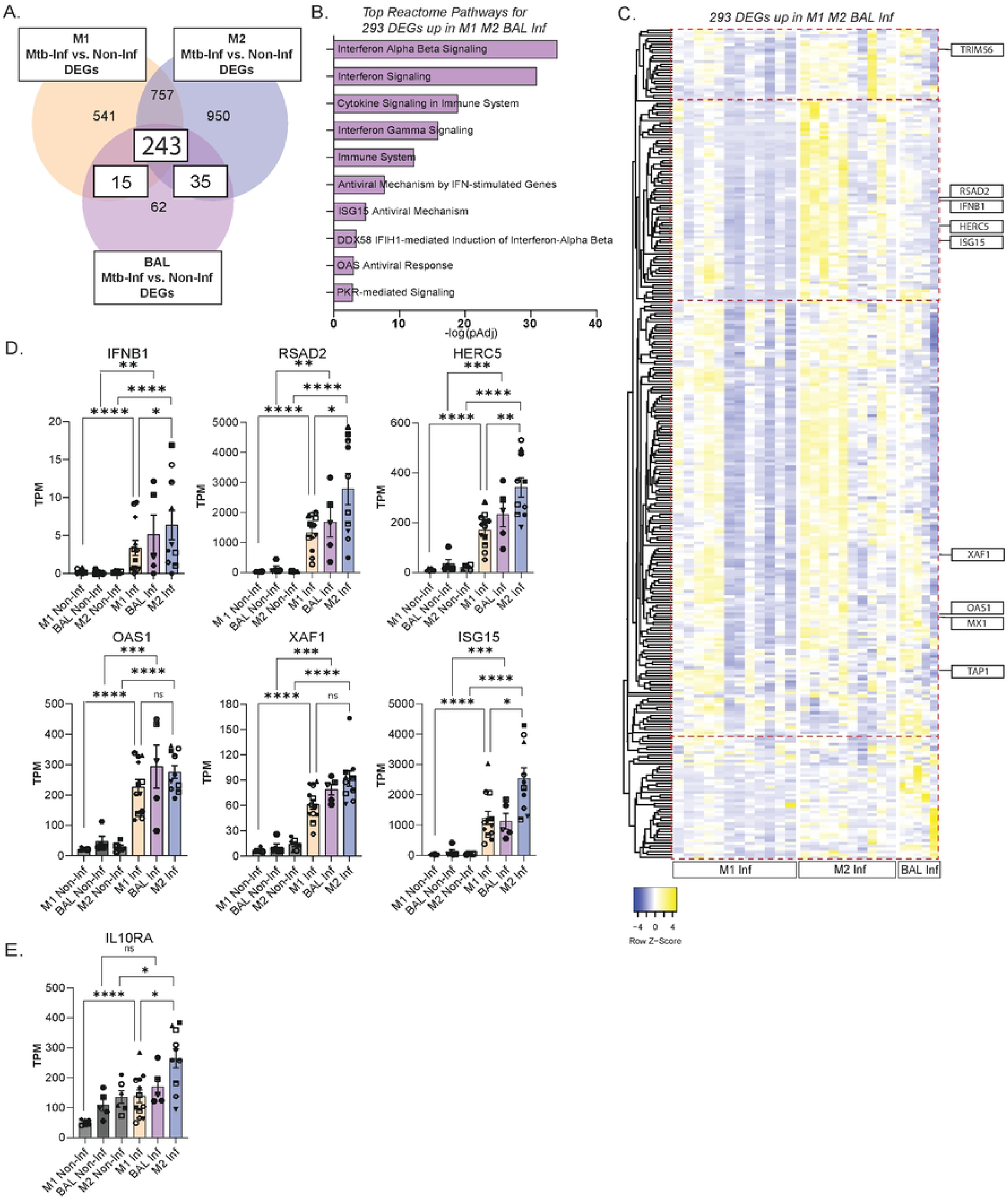
Alveolar macrophages show enrichment of key type I IFN signaling genes and *IL10RA* upon Mtb infection. **(A)** Venn diagrams showing genes upregulated in response to Mtb infection in BAL macrophages, M1- and M2-like MDMs. **(B)** Bar graph of significant (adjusted) pathways (Reactome) for the 293 DEGs upregulated by Mtb infected BAL macrophages, M1- and M2-like MDMs. Pathway enrichment significance was determined by Fisher’s exact test with Benjamini-Hochberg correction for multiple comparisons, as implemented in Enrichr. (**C**) Normalized heat map showing expression of 293 (244 genes common to all 3 subsets; 35 genes common to M2-like MDMs and BAL macrophages; 15 genes common to M1-like MDMs and BAL macrophages) DEGs upregulated in Mtb-infected BAL macrophages, M1- and M2-like MDMs. **(D)** Normalized expression counts (transcripts per kilobase-million, TPM) of selected genes from the type I Interferon pathway, each symbol represents a different donor**. (E)** Normalized expression counts (transcripts per kilobase-million, TPM) of *IL10RA*, each symbol represents a different donor. Comparisons between macrophage conditions were made using the Wald test with thresholds of log_2_ fold change > 1.0 and adjusted p value (Benjamini-Hochberg FDR) < 0.05 considered significant. * p < 0.05, ** p < 0.01, *** p < 0.001

Pathway enrichment analysis (PEA) showed Interferon Alpha/Beta signaling as the most enriched Reactome pathway **(Fig. 2B)**. Expression of the top type I IFN-stimulated genes was highest in Mtb-infected M2-like MDMs, followed by infected BAL macrophages, and lowest in infected M1-like MDMs **(Figs. 2C, 2D)**. Key genes that showed higher expression in infected BAL macrophages and M2-like MDMs over M1-like MDMs included *IFNB1, RSAD2,* and *HERC5* **(Fig. 2C, 2D)**. Importantly, not all type I IFN-related genes showed this pattern. Expression of *XAF1* and *OAS1* was similar among infected BAL macrophages, M1 and M2-like MDMs, whereas *ISG15* expression was shared by infected M1-like MDMs and BAL macrophages. These differences in type I IFN related gene expression suggested differential regulation between infected macrophage subsets.

HERC5, a HECT (homologous to the E6-AP C terminus) E3 ubiquitin ligase, is the sole human enzyme that catalyzes ISG15 attachment to host proteins, a process termed ISGylation [45,46]. ISG15 has been associated with exacerbated TB disease and was shown to be upregulated in Mtb-infected human monocytes, where it induced IL-10 secretion [39]. RSAD2 (Viperin) has been shown to inhibit IFN-γ signaling and promote lipid peroxidation in Mtb-infected murine macrophages [47,48], and is a key type I IFN induced gene. From the IL-10 pathway, *IL10RA* expression was highest in M2-like infected MDMs, followed by infected BAL macrophages, and lowest in M1-like infected MDMs **(Fig. 2E)**. These data indicate that lung resident AMs are capable of responding to IL-10 in the environment via *IL10RA* expression, and that infected M2-like MDMs and lung-resident AMs express a dominant type I IFN signaling response that involves the greater expression of *RSAD2* and *HERC5*, but not *XAF1* and *OAS1*.

### IL-10 Signaling and type I IFN pathways are enriched in *Mycobacterium tuberculosis* infected M2-like macrophages

While M2-like MDMs do not efficiently activate memory CD4+ T cells upon Mtb infection, we previously found that they successfully elicit CD4+ T cell responses when treated with Mtb lysate or γ-irradiated H37Rv (Irrad. Mtb) [9]. Therefore, we next sought to investigate the genes and pathways unique to Mtb infection of M2-like MDMs that could account for the reduced CD4+ T cell activation in response to infection. We performed bulk RNA sequencing on both M1 and M2-like MDMs 24 h after infection with Mtb (Mtb-infected), or after treatment with either γ-irradiated H37Rv (Irrad. Mtb) or media only (non-infected). Consistent with previous reports, we confirmed a large number of differentially expressed genes (DEGs) between non-infected M1 and M2-like MDMs [49,50], of which 773 remained differentially expressed after Mtb infection, while 319 DEGs were observed only with Mtb infection **(Fig. 3A)**. The expression of another 842 genes was different at baseline but changed in response to infection to become equivalent to infected M1-like MDMs **(Fig. 3B, Supplemental Fig. 1A, Extended Data 1 & 2)**. We stratified the genes from these 3 groups to focus on those with increased expression in infected M2-like MDMs, yielding a total of 1,061 genes **(Fig. 3B)**. PEA of this gene set revealed that IL-10 signaling and Interferon Alpha Beta Signaling were among the top 10 enriched Reactome pathways **(Supplemental Fig. 1B)**. Together, these results indicate type I IFN and IL-10 signaling pathways play a dominant role in the response of M2-like macrophages to infection with Mtb.

**Figure 3.**
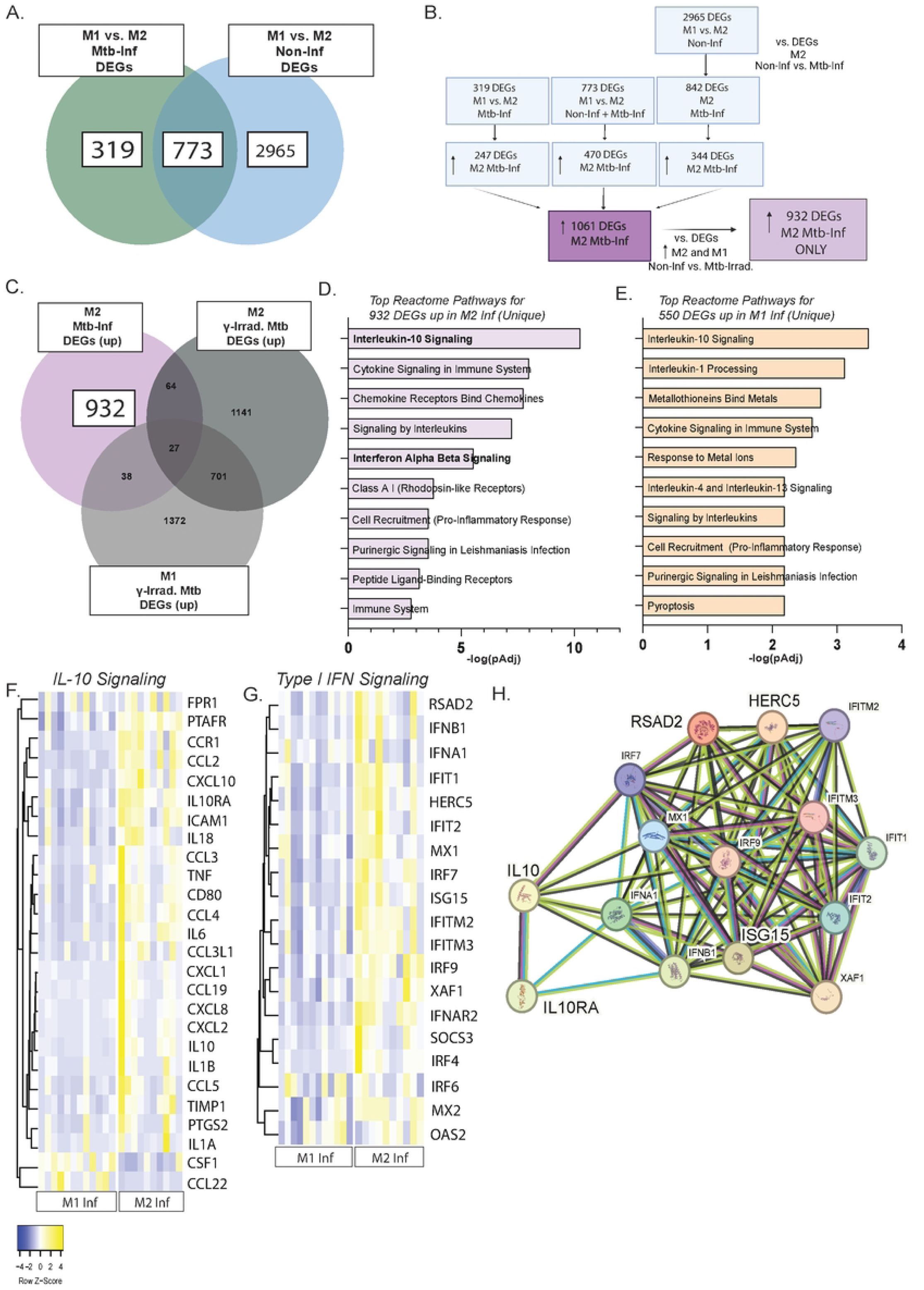
IL-10 signaling and type I IFN pathways are enriched in *Mycobacterium tuberculosis*-infected M2-like macrophages. **(A)** Venn diagram depicting DEGs between non-infected M1 and M2-like macrophages (blue), and after Mtb infection (green). **(B)** Flowchart indicating the analysis of DEGs unique to Mtb-infected M2-like macrophages, created in BioRender. **(C)** Venn diagrams depicting DEGs unique to Mtb-infected M2-like macrophages (purple) after removing upregulated DEGs in common with M1 and M2-like macrophages after Irrad. Mtb (compared to non-infected, grey). **(D)** Significant (adjusted) pathways (Reactome) of 932 genes upregulated by M2-like macrophages after Mtb infection. **(E)** Significant (adjusted) pathways (Reactome) from 550 genes upregulated by M1-like macrophages after Mtb infection. Pathway enrichment significance was determined by Fisher’s exact test with Benjamini-Hochberg correction for multiple comparisons, as implemented in Enrichr. (**F)** Normalized heat maps of IL-10 signaling pathway genes and (**G)** Type I IFN pathway genes identified in M1 and M2-like infected MDMs. **(H)** Protein-protein interaction (PPI) network generated using String DB showing evidence for interconnectedness between curated IL-10 signaling and type I IFN proteins. Line thickness indicates confidence score for the interaction. * p < 0.05; ** p < 0.01; *** p < 0.001.

Since memory CD4+ T cells were efficiently activated in response to M2-like MDMs treated with Irrad. Mtb [9], this critical control allowed us to select genes unique to Mtb-infected M2-like macrophages that could underlie impaired T cell activation. We compared the 1,061 genes upregulated in Mtb-infected M2-like MDMs with those upregulated after treatment with Irrad. Mtb relative to non-infected M1 or M2-like MDMs, revealing 932 genes unique to infected M2-like macrophages **(Fig. 3C)**. This gene set recapitulated the preferential enrichment of type I IFN and IL-10 signaling pathways in Mtb-infected M2-like MDMs **(Fig. 3D)**. A parallel analysis identified 550 genes uniquely upregulated by Mtb-infected M1-like MDMs **(Supplemental Fig. 1C, 1D)**, demonstrating IL-10 but not type I IFN signaling among the differentially expressed pathways **(Fig. 3E)**. Whereas 20 of the 22 DEGs from the IL-10 signaling pathway were expressed in infected M2-like MDMs, only 2 genes, *CSF1* and *CCL22*, drove the IL-10 signaling pathway in infected M1-like macrophages, reflecting functionally distinct responses **(Fig. 3F)**.

Type I IFN pathway genes also showed a notable dichotomy. While *SOCS3, IRF4, IRF6, MX2, XAF1*, and *OAS2* showed similar expression in both macrophage subsets, *RSAD2, HERC5, IRF7, ISG15*, *IFIT1*, *IFIT2, and IFNB1* were preferentially expressed by Mtb-infected M2-like MDMs **(Fig. 3G)**. Furthermore, the STRING protein interaction database [51] indicated strong evidence of interaction between the IL-10 signaling and type I IFN pathways **(Fig. 3H)**.

Interestingly, lower concentrations of IFN-β were detected in the supernatants of M2 compared to M1-like MDMs **(Supplemental Fig. 1E)**, whereas increased surface levels of IFNAR1 were observed **(Supplemental Fig. 1F)**, suggesting that IFN-β production is subject to enhanced negative feedback in M2-like MDMs, potentially mediated by SOCS3 [52,53]. Therefore, we focused on *RSAD2, HERC5*, and *IL10RA* as genes reliably unique to Mtb-infected M2-like MDMs. Digital PCR valdidated significant preferential expression by infected M2-like MDMs of *IL10RA* and *HERC5* with a trend toward greater expression of *RSAD2* **(Supplemental Fig. 1G, 1H)**. These data establish that IL-10 signaling and *HERC5* are preferentially enriched in Mtb-infected M2-like macrophages.

### Mtb infection of M2-like macrophages impairs MHC-II trafficking

Since both HERC5 and IL-10R signaling have been shown to impact MHC-II trafficking, we next focused on MHC-II localization in Mtb-infected macrophages [35,54]. ISGylation by HERC5 was previously shown to negatively regulate endosomal sorting complexes required for transport (ESCRT) components such as ALIX and CHMP [55], which play key roles in multivesicular body (MVB) biogenesis [54,56], and MHC-II trafficking [56]. *AP1M2* and *CHMP4B*, members of the AP-1 complex involved in MHC-II trafficking and intra-luminal vesicle formation [57,58], were found to be upregulated by infected M1 but not M2-like MDMs **(Extended Data 2)**. Importantly, the ESCRT pathway has been shown to facilitate antigen presentation to CD4+ T cells by Mtb-infected macrophages [59]. Paradoxically, M2-like macrophages exhibit high surface MHC-II expression compared to M1-like macrophages despite not efficiently activating CD4+ T cells upon infection with Mtb [9]. Therefore, we reasoned that MHC-II trafficking dynamics may be more informative than static surface staining.

To test the hypothesis that MHC-II trafficking to the plasma membrane differs between Mtb-infected M1 and M2-like macrophages, we performed an MHC-II blockade and re-staining assay with blocking and fluorescence-conjugated antibodies to compare Mtb-infected M1 and M2-like MDMs **(Fig. 4A)**. We blocked HLA-DR using saturating concentrations of unconjugated anti-HLA-DR blocking mAb (clone L243) prior to infection with GFP-expressing H37Rv (Mtb-GFP). At 24 h post infection, macrophages were then re-stained with a PE-conjugated version of the same anti-HLA-DR mAb clone (PE-L243), and also with a BV421-conjugated alternative anti-HLA-DR mAb clone, LN3 (BV421-LN3), which binds a distinct HLA-DR epitope **(Fig. 4A)**. Fluorescence microscopy revealed differences in anti-HLA-DR re-staining between M1 and M2-like MDMs. Whereas both subsets stained positively for the alternative anti-HLA-DR mAb clone (BV421-LN3), significantly fewer M2-like infected macrophages showed surface re-staining using the same clone (L243) used for HLA-DR blockade 24 h earlier **(Figs. 4B, 4C, and Supplemental Fig. 2A)**. These findings indicate that Mtb-infected M2-like macrophages show impaired trafficking of MHC-II to the plasma membrane compared to infected M1-like macrophages.

**Figure 4.**
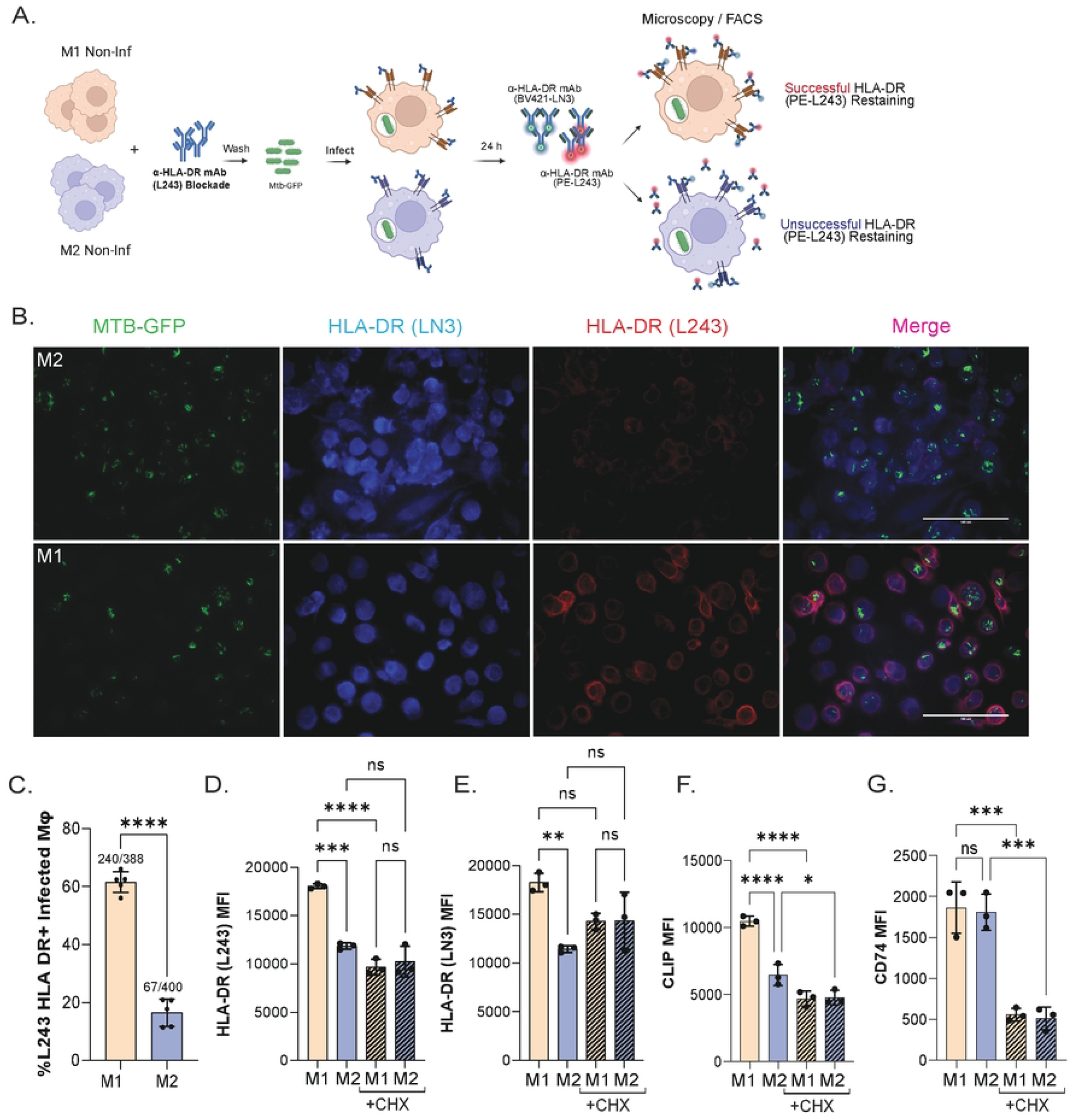
Mtb infection of M2-like macrophages impairs MHC-II trafficking. **(A)** Schematic of experimental workflow of α-MHC-II mAb blockade and re-staining assay to detect differences in MHC-II trafficking by microscopy or flow cytometry; created in BioRender. **(B)** Fluorescence microscopy of M1 and M2-like MDMs treated with anti-HLA-DR mAb blockade (clone L243), infected with Mtb-GFP showing surface anti-HLA-DR staining (BV421-LN3) and re-stained with HLA-DR (PE-L243) 24 h post infection. Images taken at 40x magnification. **(C)** Bar graph depicting the percentage of infected macrophages showing positive PE-HLA-DR (L243) re-staining, representative of 3 independent experiments (3 individuals). Significance was determined by Student’s t-test and 5-7 independent fields of view were utilized**. (D)** Bar graph showing MFI (mean + SD) of post-infection re-staining with anti-HLA-DR (PE-243) in triplicate, representative of 5 independent experiments (3 individuals) with and without treatment with cycloheximide (CHX). **(E)** Bar graph showing MFI (mean + SD) of post-infection total surface HLA-DR (BV421-LN3) in triplicate, representative of 5 independent experiments (3 individuals) with and without treatment with cycloheximide (CHX). **(F)** Bar graph showing MFI (mean + SD) of post-infection HLA-DR-bound surface CLIP in triplicate, with and without cycloheximide (CHX) representative of 2 independent experiments (2 individuals). **(G)** Bar graph showing MFI (mean + SD) of post-infection total CD74 in triplicate, with and without cycloheximide (CHX) representative of 2 independent experiments (2 individuals). Significance was determined by one-way ANOVA with Sidak’s post-test with correction for multiple comparisons. ∗∗p < 0.01; ∗∗∗∗p < 0.0001; ns, not significant.

To determine whether the impaired surface receptor trafficking in infected M2-like macrophages was specific to MHC-II, we performed a parallel assay for MHC-I trafficking. Macrophages were blocked with a saturating concentration of anti-HLA-A/B/C mAb (clone W6/32), infected with Mtb-GFP, and stained with the same anti-MHC-I mAb clone conjugated with PE (PE-W6/32) and another clone (G46-2.6) targeting a different anti-MHC-I epitope conjugated with BV421 (BV421-G46-2.6) 24 h post infection. Microscopy showed no difference between M1 and M2-like macrophages in the percentage of Mtb-infected MDMs with successful anti-HLA-A/B/C re-staining **(Supplemental Figs. 2B, 2C)**, while flow cytometry also revealed similar or slightly greater PE-W6/32 MFI in M2-like MDMs upon re-staining **(Supplemental Fig. 2C)**. These data indicate that MHC-II but not MHC-I trafficking is impaired in Mtb-infected M2-like macrophages.

We next used cycloheximide (CHX), a protein synthesis inhibitor, to determine the extent to which *de novo* MHC-II synthesis accounts for increased MHC-II trafficking to the cell surface in infected M1-like macrophages, as MHC-II surface levels depend both on new synthesis and recycling of existing molecules. After washout of Mtb-GFP, CHX was maintained in culture with infected M1 or M2-like MDMs. At 24 h post-infection, CHX treatment significantly reduced HLA-DR (L243) MFI in M1-like MDMs to levels comparable to those in M2-like MDMs **(Fig. 4D)**. By contrast, M2-like macrophages showed no significant change in PE-L243 or BV421-LN3 MFI with CHX treatment **(Figs. 4D, 4E)**. Interestingly, M1-like macrophages also showed slightly increased HLA-DR (BV421-LN3) MFI **(Fig. 4E)**, which may reflect differences in the binding kinetics of the L243 and LN3 clones. Taken together, these data indicate that MHC-II trafficking to the plasma membrane is impaired in infected M2-like macrophages and that newly trafficked HLA-DR in infected M1-like macrophages arrives at the cell surface primarily through new synthesis.

Given the reduction in MHC-II trafficking to the plasma membrane in infected M2-like macrophages, we also investigated whether other components of the antigen-processing machinery were affected. We assessed levels of CLIP (cleaved invariant chain) bound to surface HLA-DR, HLA-DM, and CD74 (invariant chain) by flow cytometry 24 h after Mtb-GFP infection of M1 and M2-like MDMs. Infected M1-like macrophages had significantly higher levels of CLIP bound to surface MHC-II **(Supplemental Fig. 2D)**, while total CD74 levels did not differ and total HLA-DM expression was lower in M1-like MDMs **(Supplemental Figs. 2E, 2F)**. As expected, CHX treatment reduced surface expression of CD74 in both macrophage subsets, while HLA-DR-bound CLIP was significantly reduced for Mtb-infected M1-like MDMs **(Figs. 4F, 4G)**. These findings further indicate that the invariant chain and HLA-DR-bound CLIP detected on the surface of infected M1-like macrophages are a product of new protein synthesis. These results also suggest that new synthesis primarily governs MHC-II molecules trafficking to the plasma membrane in Mtb-infected M1-like macrophages, whereas recycling of surface MHC-II molecules accounts for the minimal MHC-II re-staining after mAb blockade for infected M2-like macrophages.

### IL-10 neutralization and *HERC5* knockdown increase cell surface trafficking of MHC-II in Mtb-infected M2-like macrophages

Having established that MHC-II trafficking to the plasma membrane is reduced in Mtb-infected M2-like macrophages, we sought to test the hypothesis that neutralization of IL-10 and/or knockdown of *HERC5* could increase MHC-II cell surface trafficking. Using siRNA, we achieved >80% knockdown of *HERC5* in infected M2-like MDMs **(Supplemental Fig. 3A)**. *HERC5* knockdown did not significantly reduce viability or alter Mtb infection in M2-like macrophages, and surface expression of HLA-DR and CD274 showed a modest increased after *HERC5* knockdown relative to non-targeting siRNA (NT) controls **(Supplemental Figs. 3B, 3C)**.

To determine whether HERC5 and the IL-10 signaling pathway are regulated independently in infected M2-like macrophages, we measured *HERC5* expression by digital PCR in cells receiving IL-10 neutralization. Minimal decreases in *HERC5* expression were observed with IL-10 neutralization **(Supplemental Fig. 3D)**, indicating IL-10 signaling does not govern *HERC5* expression. To test the hypothesis that IL-10 neutralization or *HERC5* knockdown could improve or restore MHC-II trafficking to the plasma membrane in M2-like macrophages, we performed MHC-II blocking and re-staining experiments. Strikingly, IL-10 neutralization during and after Mtb infection of M2-like MDMs drove robust HLA-DR re-staining with PE-L243, 24 h after treatment with unconjugated anti-HLA-DR mAb blockade (also clone L243) **(Fig. 5A)**. A similar effect was observed for M2-like MDMs that received *HERC5* siRNA knockdown **(Fig. 5A)**. HLA-DR re-staining for infected M2-like MDMs treated with either IL-10 neutralization or *HERC5* siRNA knockdown was greater than in M2-like macrophages treated with isotype control or NT siRNA **(Figs. 5A, 5B, Supplemental Fig. 3E)**. Flow cytometry confirmed greater (PE-L243) HLA-DR re-staining observed with *HERC5* knockdown and IL-10 neutralization compared to controls, but similar to marginally elevated HLA-DR staining with the alternative anti-HLA-DR mAb clone (BV421-LN3) with IL-10 neutralization and *HERC5* siRNA knockdown, respectively **(Figs. 5C, 5D)**. These results indicate that IL-10 signaling and HERC5 contribute to impaired MHC-II trafficking in Mtb-infected M2-like macrophages, and that interrupting these pathways restores MHC-II trafficking and could improve antigen presentation to CD4+ T cells.

**Figure 5.**
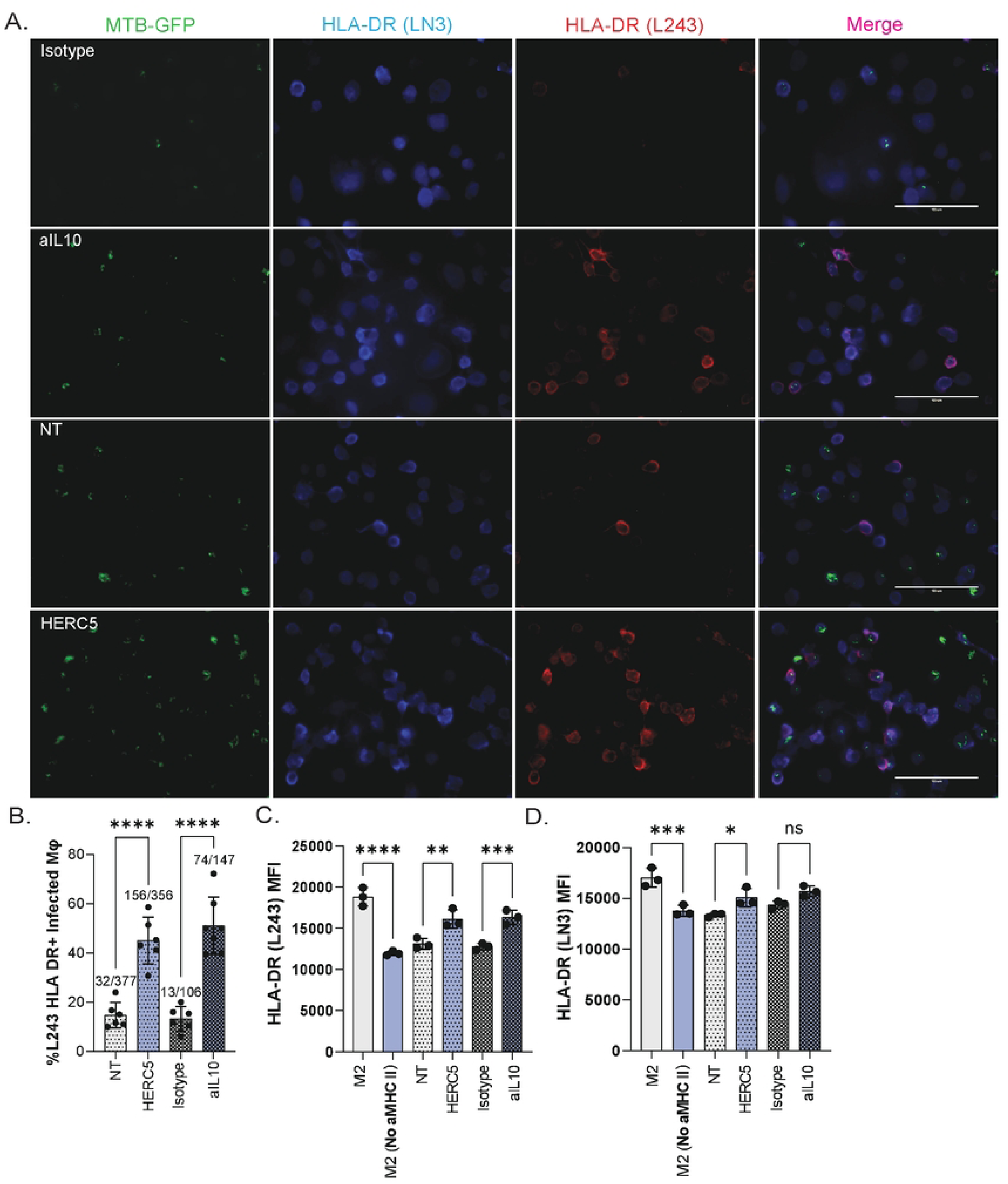
IL-10 neutralization and *HERC5* knockdown increase cell surface trafficking of MHC-II in Mtb-infected M2-like macrophages. **(A)** Fluorescence microscopy of Mtb-infected M1 and M2-like MDMs showing total surface anti-HLA-DR staining (BV421-LN3) and re-stained with HLA-DR (PE-L243) 24 h after Mtb-GFP infection of M2-like MDMs treated with an isotype mAb control (first panel), anti-IL10 neutralizing mAb (second), non-targeting siRNA control (third), and *HERC5* siRNA (fourth). Images taken at 40x magnification. (**B)** Bar graph of the percentage (mean ± SD) of infected macrophages showing positive HLA-DR (PE-L243) re-staining, representative of 3 independent experiments (3 individuals). **(C)** Bar graph showing the MFI (mean ± SD) of post-infection re-staining with HLA-DR (PE-L243) in triplicate, representative of 4 independent experiments (3 individuals) for infected M2-like MDMs under each condition. **(D)** Bar graph showing the MFI (mean + SD) of post-infection total HLA-DR (BV421-LN3) in triplicate, representative of 4 independent experiments (3 individuals) for infected M2-like cells under different conditions. Significance was determined using one-way ANOVA with Sidak’s post-test corrected for multiple comparisons. ∗∗ p < 0.01; ∗∗∗∗p < 0.0001; ns, not significant.

### IL-10 neutralization and HERC5 knockdown in Mtb-infected M2-like macrophages augments activation of memory CD4+ T cells

Since IL-10 neutralization and *HERC5* knockdown restored MHC-II trafficking to the plasma membrane of infected M2-like MDMs, we next sought to test the hypothesis that these interventions would enhance T cell activation. At 24 h post infection, memory (CD45RA^Lo^) CD4+

T cells from healthy individuals with LTBI were isolated by immunomagnetic negative selection and co-cultured with autologous, Mtb-infected MDMs for 16 h. A significantly higher proportion of CD4+ T cells were activated in response to infected M2-like MDMs treated with *HERC5* siRNA knockdown (0.38% [0.2067–0.7033]; median [interquartile range (IQR)], 11 participants), IL-10 neutralization (0.583% [0.356–0.807]), or both (0.59% [0.263 –0.847]), as measured by co-expression of CD69 and CD40L, relative to non-targeting siRNA control (0.245% [0.090–0.443]) **(Figs. 6A, 6B)**. CD69 and CD40L co-expression was also reduced with anti-MHC-II (HLA-DR/DQ/DP) mAb blockade (0.291% [0.136—0.411), confirming the augmented T cell activation was TCR-pMHC-II-dependent. Co-expression of CD69 and IFN-γ was highest for infected M2-like MDMs treated with IL-10 neutralization in combination with HERC5 knockdown (0.211% [0.067—0.238]; median [IQR], 10 participants), IL-10 neutralization (0.153%, [0.104—0.3617]), and for *HERC5* knockdown (0.09 [0.043—0.195]), relative to NT controls 0.049 [0.020—0.118]) **(Figs. 6C, 6D)**. These results link the restoration of MHC-II trafficking by IL-10 neutralization and *HERC5* knockdown to augmented CD4+ T cell responses Mtb-infected M2-like macrophages.

**Figure 6.**
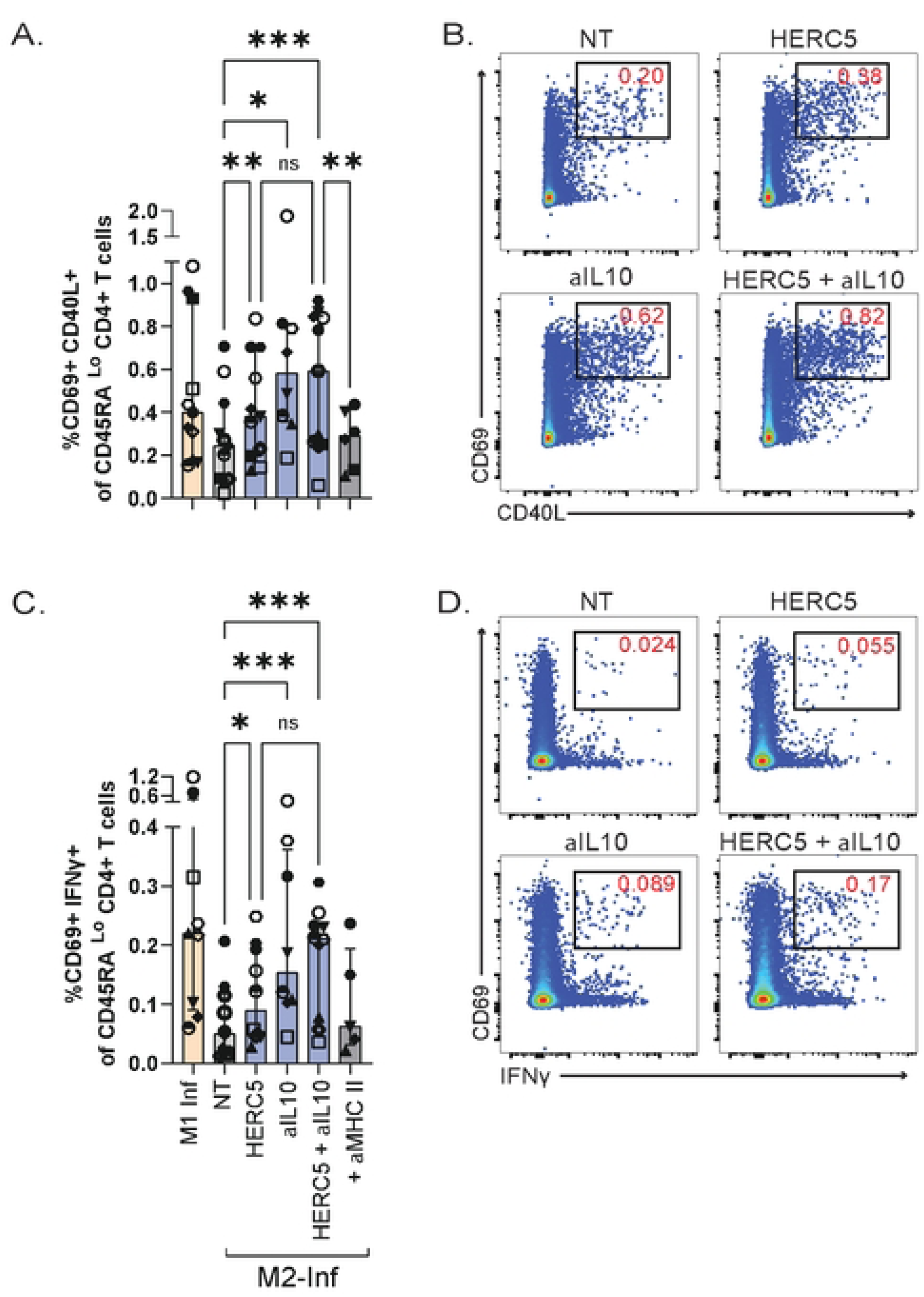
IL-10 neutralization and *HERC5* knockdown in Mtb-infected M2-like macrophages augments activation of memory CD4+ T cells. **(A)** Summary Bar graphs and **(B)** representative flow cytometry plots comparing median (and IQR) co-expression of CD69 and CD40L gated on CD45RA^Lo^ (memory) CD4^+^ T cells after 16-18 h co-culture with Mtb infected macrophages in each condition. (**C**) Summary Bar graphs and **(D)** representative flow cytometry plots comparing median (and IQR) co-expression of CD69 and IFN-ү for memory CD4+ T cells in response to Mtb-infected macrophages in the same conditions. Data represent 10-12 individuals from 4-5 independent experiments. Significance was determined by paired mixed effects analysis of variance and Sidak’s post test, corrected for multiple comparisons. ∗ p < 0.05; ∗∗ p < 0.01; ∗∗∗p < 0.001; ns, not significant.

## Discussion

Macrophages, the cellular niche for Mtb infection, can restrict bacterial growth or provide an environment permissive to Mtb persistence. In this study, we expand on our previous work demonstrating the differential capacity of macrophage subsets to activate CD4+ T cells upon Mtb infection. We find that compared to M1-like MDMs, M2-like macrophages exhibit greater type I IFN and IL-10 signaling in response to Mtb infection. Similar to M2-like MDMs, AMs expressed key type I IFN genes including *HERC5* and *RSAD2,* high levels of *IL10RA,* and failed to efficiently activate memory CD4+ T cells. We further show that MHC-II trafficking to the plasma membrane is uniquely impaired in infected M2-like MDMs, and is restored with IL-10 neutralization or *HERC5* knockdown, which leads to augmented memory CD4+ T cell activation. Together, these data link increased IL-10 signaling and *HERC5* expression in Mtb-infected M2-like macrophages to impaired antigen presentation and dampened CD4+ T cell activation.

Constitutive IL-10 signaling in the lung is likely critical for maintaining gas exchange, but could be exploited by Mtb during infection of macrophages expressing IL-10R [9,35,37,60–62]. To examine the link between IL-10 neutralization and enhanced T cell activation, we focused on antigen presentation. We found MHC-II trafficking to the plasma membrane to be impaired among Mtb-infected M2-like macrophages in the presence of IL-10 signaling. Compared to M1-like macrophages, M2-like MDMs secreted high levels of IL-10 at baseline and after Mtb infection. Non-infected M2-like MDMs showed expression of *IL10RA* compared to M1-like MDMs. Infected M2-like MDMs showed the greatest expression, followed by infected AMs from BAL samples. These results indicate that AMs and M2-like MDMs demonstrate increased sensitivity to IL-10 signaling relative to M1-like MDMs, and that IL-10 signaling can facilitate a state of restricted MHC-II exchange between the cell surface and the phagosome or other intracellular compartments during infection. The preferential expression of type I IFN genes in infected M2-like macrophages, previously associated with phagosomal membrane damage and permeability [63,64], also provide a link to restricted MHC-II exchange. We speculate this could result either from reduced docking of MHC-II at damaged phagosomal membranes or increased retention of MHC-II within these membranes, perhaps as a result of autophagy [65]. Either mechanism would reduce cell surface MHC-II levels, diminish presentation of Mtb-derived peptides on the cell surface, and, consequently, impaired CD4+ T cell activation. Since IL-10 signaling has also been linked to reduced phagosome maturation [33], Mtb pathogenesis in M2-like macrophages is likely exacerbated by both impaired phagosomal proteolysis and hindered antigen presentation arising from diminished MHC-II loading and trafficking.

The type I IFN response depends on phagosomal membrane damage by Mtb infection, delivering bacterial DNA and secreted proteins into the cytosol [63,64], and is associated with uncontrolled bacterial growth [10,13,29]. Multiple studies have shown an interconnection between IL-10 and type I IFN pathways [37,62,66]. We observed preferential enrichment of key type I IFN genes including *HERC5* and *RSAD2* in Mtb-infected M2-like macrophages. However, other genes in the type I IFN pathway, such as *XAF1* and *OAS1*, were not enriched. The most likely explanation is that these genes are also induced by other pathways. *XAF1* has roles as a tumor suppressor gene with anti-apoptotic function [67], and can be induced by both the RANKL and RIG-I/MAVS pathways [68]. *OAS1* encodes a key enzyme involved in antiviral defense and RNAse activation, but, like *XAF1*, it is also induced by multiple signaling pathways including IFN-ү signaling and IL-6/JAK/STAT [69]. We selected *HERC5* as a knockdown target based on its role as the sole enzyme catalyzing the addition of ISG15 to host proteins, previously studied in antiviral responses [45,46,70]. Prior work has shown that HERC5 limits viral replication by preventing viral budding from infected cell membranes [70] and potentiates the type I IFN response by stabilizing cGAS [46]. HERC5 expression also promotes degradation of key ESCRT pathway components, including Tsg101 and ALIX [55]. While the ESCRT pathway is used by many viruses to exit cells during assembly, and its inhibition limits viral spread [55], ESCRT proteins also play a role in MHC-II trafficking [56–58], phagosomal membrane repair [71,72], and CD4+ T cell activation by Mtb-infected macrophages [59]. Disrupting ESCRT machinery during infection impairs control of mycobacterial growth, potentially by compromising phagosomal components required for bacterial growth restriction [73]. Phagosomal membrane damage can also promote antigen leakage into the cytosol. In one study, sustained phagosomal disrepair enhanced DC cross-presentation of bacterial antigens to CD8+ T cells [74]. Impaired phagosome maturation and decreased protein degradation likely reduce antigen availability for MHC-II loading, contributing to impaired antigen presentation to CD4+ T cells.

Similar to IL-10 neutralization, *HERC5* knockdown in M2-like macrophages also increased CD4+ T cell activation. These pathways appear to be independent as IL-10 blockade of infected M2-like MDMs only modestly decreased *HERC5* expression. IL-10 neutralization alone or in combination with *HERC5* knockdown yielded the greatest CD4+ T cell activation. This likely reflects several factors. *HERC5* knockdown is probably most effective for individuals with a greater dynamic range of *HERC5* expression in macrophages. For people who do not mount potent type I IFN responses to infection, IL-10 neutralization may more effectively drive the augmented CD4+ T cell activation. Methodological differences may also contribute, as antibody neutralization could be more complete than the 80-85% HERC5 depletion achieved by siRNA treatment, and a small-molecule inhibitor of HERC5 might therefore produce more consistent results. Furthermore, combined blockade did not show additive or synergistic improvement of MHC-II trafficking or CD4+ T cell activation, suggesting overlapping effects on antigen presentation to CD4+ T cells. If HERC5-mediated inhibition of the ESCRT pathway limits phagosomal membrane repair during Mtb infection, host-directed therapies targeting HERC5 could restore phagosomal integrity, improve MHC-II trafficking, and enable efficient Mtb antigen loading. A recent study also showed that glycolytic enzymes are major substrates for HERC5-mediated ISGylation, which can promote their degradation, shifting macrophages toward oxidative phosphorylation [75]. This metabolic shift is supported by studies showing that type I IFN signaling rewires macrophages to favor oxidative phosphorylation (OXPHOS) and fatty acid oxidation (FAO) over glycolysis in multiple immune contexts [76,77] including during mycobacterial disease [78]. OXPHOS and FAO are metabolic programs typically associated with anti-inflammatory macrophages, whereas glycolysis is suggestive of more immune-activated APCs [79–82]. Furthermore, glycolytic enzymes such as Aldolase A have been linked to phagosomal maturation. Aldolase A facilitates V-ATPase assembly at phagosomes in *Salmonella typhimurium*-infected macrophages, enabling phagosomal acidification and bacterial restriction [83]. A glycolysis-to-oxidative phosphorylation shift could therefore impair phagosomal maturation and reduce Mtb antigen loading onto MHC-II. Finally, ISG15 deficiency has been associated with increased susceptibility to mycobacterial disease [84]. However, its protective function is attributed to free ISG15 acting as a cytokine to promote IFN-γ signaling, independent of its conjugation activity. Inhibition of HERC5 could therefore enhance T cell activation by increasing the relative abundance of free ISG15 in Mtb-infected M2-like macrophages.

Previous studies have implicated both IL-10 signaling and type I IFN responses as detrimental to protection against TB [10,13,28–34]. However, a clear mechanism connecting these pathways to T cell recognition of Mtb-infected human macrophages has not been established. Our work identifies the roles of IL-10 signaling and *HERC5* in impairing MHC-II trafficking and thereby reducing antigen presentation by infected M2-like macrophages to CD4+ T cells. Inhibition of these pathways restores MHC-II trafficking and enables T cell activation, possibly through reduced phagosomal membrane damage facilitated by ESCRT-mediated repair, improved antigen retention in the phagosome, and efficient loading onto MHC-II or increased glycolytic flux and enhanced phagosomal maturation enabling proteolysis and presentation of Mtb antigens. Targeted inhibition of IL-10 signaling and HERC5 in infected M2-like macrophages could represent host-directed therapies capable of reprogramming anti-inflammatory macrophages to activate CD4+ T cells, suppressing infection and preventing or treating active TB.

### Limitations of the study

To our knowledge, this study is the first to demonstrate a direct link between type I IFN signaling and impaired MHC-II trafficking during Mtb infection of human macrophages. Nevertheless, several limitations should be noted. First, although Mtb-infected M2-like MDMs represent many aspects of infected macrophages in the lungs, our model does not fully recapitulate the spectrum of diverse macrophage phenotypes found *in vivo.* Our *ex vivo* macrophage infection model represents extreme but useful M1 and M2 polarized phenotypes. Aside from limited macrophage phenotypes, we were similarly limited in our ability to examine the roles of type I IFN pathway genes. We focused on *HERC5* based on its strong and consistent expression in infected M2-like macrophages, but other type I IFN signaling genes likely also contribute to reduced antigen presentation to CD4+ T cells. An obvious constraint of our study is the small sample size. We were limited by the scarce availability of human BAL samples for research, particularly from individuals with LTBI, which hampers the applicability of our findings across populations. Donor-to-donor variability likely contributed to variation in the magnitude of responses to IL-10 neutralization and *HERC5* knockdown in both MHC-II trafficking and T cell activation experiments. While we observed consistent CD4+ T cell activation with either intervention, it is possible that with a larger sample size, greater MHC-II trafficking and CD4+ T cell activation responses could have been observed with HERC5 knockdown, or in combination with IL-10 neutralization. Finally, since our experiments focused exclusively on infected macrophage and CD4+ T cell interactions, they do not capture the full complexity of cellular interactions likely occurring *in vivo*. How interactions with other cell types influence infected macrophages and T cells remains an open question. Despite these limitations, this work provides mechanistic insight into how IL-10 signaling and HERC5 could be modulated to enhance CD4+ T cell responses to infected macrophages to treat or prevent active TB.

## Materials and Methods

### Human participants

Twelve healthy participants ages 25-63 (eight male, four female) representing African American, Asian, White, and Latinx ethnicities were prospectively enrolled as volunteers for blood draws between September 1^st^ 2023 and September 30^th^ 2025. Participants were either referred from the MetroHealth TB clinic in Cleveland, OH, based on a positive tuberculin skin test (TST), or volunteered from University Hospitals or the Case Western Reserve University campus in response to flyers, based on a self-identified history of LTBI. For all individuals, blood was drawn only after written informed consent was obtained and the risks and benefits of the study were explained. De-identified sample IDs were used for all samples and no personally identifying information was used for the purpose of this study. LTBI status was determined by tuberculin skin test (TST), IFN-γ release assay (IGRA), or both, and confirmed using QuantiFERON-TB Gold Plus (Qiagen, Hilden, Germany). Samples from one participant were excluded from analysis due to low cell viability. Seven participants previously received isoniazid or rifampin at least 3 years prior to participating. Six individuals previously received the BCG vaccine as infants.

Cryopreserved BAL and PBMC samples from 10 additional healthy volunteers, ages 21-47 (six male, four female) were utilized between 10/1/2024 and 8/30/2025 to study pulmonary immune cells in the context of Mtb infection. Authors performing experiments and analysis for this study did not have access to information that could personally identify individual participants during or after experiments or data collection; only de-identified sample IDs were used. Participants volunteered from the Case Western Reserve University campus for research bronchoscopy, including five individuals with LTBI using the same criteria, and five were IGRA-negative. For all individuals, research bronchoscopy and blood draws were performed only after written informed consent was obtained and the risks and benefits of the study were explained. Male and female participants were eligible for research bronchoscopy based on age (18–50), non-smoking status, and lack of other significant medical problems including asthma or other chronic respiratory disease, cardiac disease, or ongoing use of systemic immunosuppressive agents for any reason. Samples from one participant were excluded from analysis due to low cell viability.

For macrophage studies, seven leukapheresis products (leukopaks) from healthy individuals ages 26-53 (5 male, 2 female) were purchased from AllCells (Alameda, CA, USA); these participants were separate from the LTBI participants. All procedures involving human subjects were approved by the Institutional Review Boards of University Hospitals Cleveland Medical Center or MetroHealth, Cleveland, OH, USA, and written informed consent was obtained from all participants.

### Generation of monocyte-derived macrophages

Following blood draw, diluted whole blood was underlaid with Ficoll-Paque PLUS (GE Healthcare, Chicago) using a Pasteur pipette. After centrifugation, the PBMC layer was aspirated with a transfer pipette. The PBMCs were then washed with Ca^++^- and Mg^++^-free and pyrogen-free sterile phosphate-buffered saline (hereafter PBS; Corning, NY) and counted by hemocytometer after trypan blue (Life Technologies) exclusion of dead cells. CD14^+^ monocytes were then isolated from PBMCs by positive immunomagnetic selection using anti-human CD14 microbeads (Miltenyi Biotec, Germany), per the manufacturer’s instructions. After CD14 selection, both the positive and negative fractions were counted and separately cryopreserved in freezing medium (10% DMSO, 90% FBS) in liquid nitrogen.

For T cell experiments, CD14+ monocytes were thawed and rested overnight in complete RPMI 1640 medium (cRPMI) containing 10% FBS, 2 mM L-glutamine, Na-pyruvate, nonessential amino acids, 0.01 M HEPES buffer, 2-mercaptoethanol, and 0.005 M NaOH (Gibco), as described previously [43]. The next morning, CD14^+^ cells were plated in 24-well plates (250,000 cells/well) (Corning, NY) and differentiated to macrophages using GM-CSF or M-CSF (50 ng/ml) (PeproTech, East Windsor, NJ, USA). Half the medium in each well was replaced at day 3 with GM-CSF- or M-CSF-containing cRPMI. After 6 days, macrophages were ready for Mtb infection and medium was exchanged for fresh cRPMI immediately prior to infection, as described previously [43].

### Bronchoalveolar lavage (BAL) cell culture

BAL samples were thawed and washed in cRPMI supplemented with penicillin/streptomycin (Pen/Strep) and Fungizone (Gibco). The sample was enriched for live cells using a Dead Cell removal kit (Miltenyi Biotec) following manufacturer instructions, as previously described[9,85]. BAL cells were plated in either 24 well (250,000/well) or 96-well flat-bottom tissue culture-treated plates (50,000/well) in cRPMI supplemented with Pen/Strep (Cytiva, MA, USA) and Fungizone (Gibco). Macrophages were allowed to adhere over 3-4 hours. Non-adherent cells were removed and cultured separately. Adherent macrophages were infected with Mtb strain H37Rv at an MOI of 4–5. After 4 h, extracellular bacteria were washed out and macrophages were incubated overnight, as previously described [9]. Approximately 24 h later, non-adherent cells or total CD4+ T cells from PBMCs were added to BAL macrophages or MDMs. After 16-18 h, non-adherent T cells were harvested, stained with fluorescence-conjugated antibodies, and prepared for analysis by flow cytometry, described below.

### Microbe strains, Bacterial Culture and Infection

One mL aliquots of *Mycobacterium tuberculosis* strain H37Rv (NR-13648, BEI Resources) or H37Rv-GFP (kindly provided by Dr. Bryan Bryson, MIT Broad Institute, Cambridge, MA, USA) were thawed, diluted in 9 mL of Difco Middlebrook 7H9 medium, and expanded to mid-log phase over 6 days, as described previously [43]. After reaching an optical density at 600 nm (OD600) of 0.3-0.8, cultures were washed with cRPMI and filtered through a 5-µm Millex syringe-driven filter unit (Millipore) to generate a single-cell bacterial suspension. Bacterial density was determined by OD600 of filtered bacteria resuspended in cRPMI, and bacteria were added to macrophages at a target multiplicity of infection (MOI) of 1-3 bacteria per macrophage, as described previously [9,43]. After 4 h, wells were washed to remove extracellular bacteria and incubated overnight in fresh cRPMI.

### Sample processing for Bulk RNA sequencing

On day 6, M1-like MDMs, and M2-like MDMs were infected with H37Rv or H37Rv-GFP, or treated with gamma-irradiated H37Rv (10 mg/mL), or left uninfected (cRPMI only). Adherent AMs plated from cryopreserved BAL samples were also infected (H37Rv) or left uninfected. At 24 h post infection, 1.0 mL TRIzol (Invitrogen) was added to each well of adherent macrophages in 24-well plates for 5 min at room temperature and homogenates were stored at −80°C until RNA isolation. Total RNA was isolated by thawing cell lysates, adding BAN phase separation reagent (Thermo) to the samples, and transferring them to phase separator tubes (Thermo). The aqueous phase containing RNA was then added to Qiagen RNAeasy columns and RNA was isolated according to the manufacturer’s protocol (Qiagen, Hilden, Germany). RNA was quantified by Nanodrop and Qubit 4 Fluorometer (Invitrogen) and at least 100 ng of RNA for each sample was submitted to GENEWIZ® NGS Services from Azenta Life Sciences (South Plainfield, NJ, USA) for bulk RNA sequencing. Quality control was performed using a Tape Station (Agilent, Santa Clara, CA, USA) and samples with a RIN ≥ 7 or DNV ≥ 200 were used. Samples were treated with TURBO DNase (Thermo), and rRNA was depleted using QIAGEN FastSelect rRNA Bacteria + HMR kit (Qiagen, Germantown, MD, USA) according to manufacturer’s protocol. RNA sequencing libraries were constructed with NEBNext Ultra II RNA Library Preparation Kit for Illumina by following the manufacturer’s guidelines. Briefly, enriched RNA was fragmented for 15 min at 94°C, and first and second strand cDNA were synthesized. cDNA fragments were end-repaired and adenylated at the 3’ end, universal adaptors ligated to cDNA fragments, followed by index addition and library enrichment with limited cycle PCR according to the manufacturer’s instructions. Sequencing libraries were validated using the Agilent Tapestation 4200 (Agilent Technologies, Palo Alto, CA, USA), and quantified using Qubit 4 Fluorometer (Thermo) and quantitative PCR (KAPA Biosystems, Wilmington, MA, USA). Sequencing libraries were multiplexed and clustered on the flowcell. After clustering, the flowcell was loaded on the Illumina NovaSeq instrument according to the manufacturer’s instructions. Samples were sequenced using a 2×150 Paired-End (PE) configuration. Raw sequence data (.bcl files) generated from the NovaSeq was converted into fastq files and demultiplexed using Illumina bcl2fastq 2.20 software. One mismatch was allowed for index sequence identification.

### Differential Gene Expression Analysis

After investigating the quality of the raw data, sequencing reads were trimmed to remove possible adaptor sequences and nucleotides with poor quality using Trimmomatic v.0.36. Trimmed reads were mapped to the GRCh38.p14 reference human genome available on ENSEMBL using STAR aligner v.2.5.2b. BAM files were generated following this step. Following alignment, unique gene hit counts were quantified using featureCounts from the Subread package v.1.5.2 to generate a gene count expression matrix across all macrophage samples. Only unique reads that fell within exon regions were counted. After extraction of gene hit counts, the gene hit counts table was used for downstream differential expression analysis. Differential gene expression analysis comparing infected and non-infected pairs of BAL macrophages/AMs, M1, and M2-like MDMs was performed using DESeq2 [86], using the Wald test with thresholds of log2 fold change > 1.0 and adjusted p value (Benjamini-Hochberg FDR) < 0.05. Gene ontology analysis was performed on the statistically significant set of genes by implementing the software GeneSCF. The species GO list was used to cluster the set of genes based on their biological process and determine their statistical significance. PCA was performed using the “plotPCA” function within the DESeq2 R package. The top 500 genes, selected by highest row variance were used to generate the plot. Venn diagrams were generated using JVenn [87], and pathway enrichment analysis was performed with Enrichr [88] using the Reactome database (2024). Raw TPM values were used to create bar graphs in Prism. TPM values were normalized using Prism and used to generate heat maps using Heatmapper [89].

### siRNA Knockdown and Digital PCR Quantification

siRNA knockdown of M2-like MDMs was performed using the ON-TARGETplus siRNA smartpool (Horizon Discovery, UK) for *HERC5* and non-targeting control, and the Mirus-X2 transfection reagent (Mirus Bio, WI, USA). On day 6 post differentiation, 25 nM of *HERC5* or non-targeting siRNA pools were complexed in OPTI-MEM media (Mirus Bio) according to the manufacturer’s protocol, using 1.0 µL of Mirus-X2 instead of the recommended 1.5 µL for improved cell viability. siRNA complexes were added to M2-like MDM wells in cRPMI. At 10–12 h post-knockdown, wells were washed and fresh cRPMI was added. At 24 h post-knockdown, MDMs were infected with Mtb strains H37Rv or Mtb(H37Rv)-GFP as described above.

To validate RNA sequencing gene expression differences, and to confirm siRNA knockdown efficiency, digital PCR was performed using the AbsoluteQ system (Thermo). Briefly, at 24 h post infection TRIzol was added to MDMs as described above. For RNA isolation, the aqueous phase was separated using phase separation tubes (Thermo) and transferred to Qiagen RNAeasy columns. First-strand cDNA synthesis was performed using 50–60 ng of total RNA with the SuperScript IV VILO mastermix (Thermo) per the manufacturer’s protocol. Digital PCR was then performed on diluted or undiluted cDNA using the AbsoluteQ system (Invitrogen), according to the manufacturer’s protocol. TaqMan primer probes (Thermo) were used to quantify each gene target. Absolute counts per μL of sample were determined using QuantStudio Digital PCR Software v6.

### Immunofluorescence microscopy and image analysis

Approximately 120,000 CD14+ monocytes were plated in wells on chamber slides (Nunc) and differentiated in cRPMI for 6 days using either M-CSF or GM-CSF as indicated. For siRNA knockdown conditions, reagents were calculated based on the surface area of the individual well of the chamber slide and performed as indicated above. 24 h after siRNA treatment, 25 ug/ml of anti-HLA-DR mAb was added to wells for 20 min in cRPMI at 37 C 5% CO2. Media was removed and macrophages were infected with H37Rv-GFP at MOI 1. 24 h post infection, cells were washed with PBS, treated with Fc receptor blockade with TruStain reagent (Biolegend, San Diego, CA, USA) for 20 minutes in AutoMACS running buffer at 4C, followed by staining with anti-HLA-DR PE (L243, BD) and HLA-DR BV421 (LN3, Thermo) mAbs in AutoMACS running buffer for 20 min at 4C. Following washout with PBS, cells were fixed with 1% PFA in PBS for 1 hour prior to removal from the BL-3, according to institutional biosafety protocols. PFA was removed and Pro-Long Diamond mounting media (Thermo) was added to each sample. Samples were cured at RT for ∼24 hours and visualized. Images were acquired on the EVOS FL Auto inverted fluorescent imaging system (Thermo). Images were taken using the DAPI, GFP, and RFP light cubes (Thermo) and 20X and 40X magnifications were used. BV421-HLA-DR was imaged in the DAPI channel, H37Rv-GFP was imaged in the GFP channel, and PE-HLA-DR was imaged in the RFP channel. 4-8 random fields of view covering all 4 quadrants of each well on the slide were obtained using EVOS FL Auto software (Thermo). Raw images were processed using ImageJ (NIH, USA).

### ELISA and flow cytometry analysis of macrophages

To quantify IFNβ production, macrophage culture supernatants were removed from MDMs 24 h post Mtb infection. IFNβ ELISA (Biolegend) was performed on neat supernatants. A standard curve was generated in duplicate, and ELISA samples were analyzed using the Synergy H1 microplate reader (BioTek, Winooski, VT). For flow cytometry, macrophages were harvested from plates 24 h post-infection using incubation with Accutase cell detachment solution (Corning, NY) for 10 minutes at 37 C, as described previously [9]. Harvested macrophages were transferred to microcentrifuge tubes and centrifuged at 500 × g for 5 min at 4C. Cell pellets were resuspended in PBS and stained with fixable Live/Dead Near-IR viability dye (Invitrogen) according to the manufacturer’s instructions, followed by fluorescence-conjugated antibodies for flow cytometry. For HLA-DR blockade conditions, anti-HLA-DR mAb (clone L243, Ultra-LEAF; BioLegend) was added to macrophages at 25 µg/mL in cRPMI for approximately 20 min at 37°C, 5% CO₂. Media was then removed and cells were infected with H37Rv. For conditions that received cycloheximide (CHX) (Thermo), lyophilized cycloheximide was resuspended in 100% DMSO, diluted in cRPMI at a final concentration of 10 μM (0.001% DMSO), and sterile filtered using a 0.22 micron filter. CHX containing cRPMI was added to MDMs after washout of bacteria and kept in culture until harvest. For conditions receiving either isotype (Ultra-LEAF™ Purified Rat IgG2a, κ Isotype, Biolegend) or IL-10 neutralization (Ultra-LEAF purified anti-human IL-10 (clone JES3-19F1), Biolegend), respective antibody was added to macrophages during infection, and after washing out extracellular bacteria at a final concentration of 10 ug/ml. Prior to removal from the BSL-3, cells were fixed with 1% PFA for 1 hour according to institutional biosafety protocols. For CD74 and HLA-DM staining, macrophages were permeabilized post fixation with PFA using the Perm/Fix kit according to manufacturer instructions (BD Biosciences, Franklin Lakes, NJ) and stained with fluorescence-conjugated antibodies. Samples were acquired on a BD LSRFortessa X-20 Cell Analyzer (BD Biosciences).

Fluorescent antibodies used in flow cytometry included: BV785-conjugated anti-human CD11b (ICRF44) (RRID: AB_2563794), BV421 anti-CD210α (3F9) (RRID:AB_2728246), PE anti-HLA-ABC (W6/32) (RRID:AB_314874), BV711 anti-PD-L1 (29E.2A3) (RRID: AB_2565764) from Biolegend, and PE anti-HLA-DR (L243) (RRID:AB_396510), BV650 anti-CLIP (cerCLIP) (RRID:AB_2740898), BV421 anti-HLA-ABC (G46-2.6) (RRID:AB_2739189), PE anti-HLA-DM (MaP.DM1) (RRID:AB_396272) and BV421 anti-CD74 (LN2) (RRID:AB_2741705) from BD Biosciences, and BV421 anti-HLA-DR (LN3) (RRID:AB_2942150) from eBioscience.

### CD4 memory T cell co-culture assay

On the day of macrophage infection, aliquots of CD14-negative PBMCs were thawed and rested overnight in cRPMI. The following day, memory CD4+ T cells (CD45RAlo) were isolated by negative immunomagnetic selection using the Human Memory CD4 T Cell Isolation Kit (Miltenyi Biotec) per the manufacturer’s instructions, with AutoMACS Rinsing Solution used as the wash buffer. After selection, CD4+ T cells were added to infected macrophages at approximately a 1:1 or 2:1 ratio (250,000–500,000 T cells per 250,000 macrophages in 24-well plates). Anti-CD40 blocking antibody (clone W17212H, BioLegend) was added simultaneously with T cells at a final concentration of 0.5 µg/mL. Anti-CD40 blockade was used to prevent CD40 internalization on Mtb-infected macrophages, enabling detection of CD40L on T cells. CD4+ T cells were co-cultured with infected macrophages in cRPMI at 37°C, 5% CO₂ overnight. After 16 h, T cells were harvested by vigorous pipetting. For most experiments, the Miltenyi IFN-γ Secretion Assay was performed per the manufacturer’s instructions, followed by viability dye and surface antibody staining, and fixation with 1% PFA for 1 hour, followed by washout with AutoMACS running buffer prior to flow cytometry. Samples were acquired on the LSR Fortessa X-20 Cell Analyzer (BD Biosciences). Samples with > 70% cell viability were included in the analysis. Anti-IL-10 mAb (clone JES3-19F1) was added to certain macrophage conditions during infection, after washout of extracellular bacteria, and again at the time of T cell co-culture, at a final concentration of 10 µg/mL, and remained in culture until T cell harvest. For MHC-II blockade conditions, a cocktail of anti-HLA-DR (clone L243), anti-HLA-DP (clone B7/21), and anti-HLA-DQ (clone 1a3) antibodies was added to Mtb-infected macrophages prior to T cell addition at 25 µg/mL each. Non-targeting siRNA and MHC-II blockade conditions served as negative controls. Functional antibodies used were mAbs against: HLA-DR and IL-10, Ultra-LEAF grade (low endotoxin, azide-free; BioLegend); and HLA-DQ and HLA-DP, In Vivo GOLD grade (Leinco).

Fluorescent antibodies used in flow cytometry included: BUV395 (RRID: AB_2738770) - conjugated anti-human CD69 (FN50), BUV737 anti-CD3 (UCHT1) (RRID: AB_2870081), (), and BB515 anti-CD25 (M-A251) (RRID: AB_2739065), from BD Biosciences, and BV785 anti-CD4 (RPA-T4) (RRID: AB_2564382), APC-Fire-750 anti-CD45RA (HI100) (RRID: AB_2616715), BV711 anti-PD-L1 (29E.2A3) (RRID: AB_2565764) and PE-Dazzle-594 anti-CD40L (24-31) (RRID: AB_2566245) from Biolegend.

### Data analysis and statistics

Flow cytometry data were analyzed using FlowJo v10 (BD Biosciences, San Diego, CA), and the proportion (percentage) and/or median fluorescence intensity (MFI) of each marker were compared between groups. For macrophages, markers were compared after gating on Live/Dead−CD11b+ cells. Pairwise comparisons of normalized gene expression (TPM) between macrophage conditions were made using the Wald test with thresholds of log_2_ fold change > 1.0 and adjusted p value (Benjamini-Hochberg FDR) < 0.05. Digital PCR data were analyzed using QuantStudio Absolute Q Digital PCR Software v6. Mean mRNA concentrations from 2-3 biological replicates were compared between conditions using one-way ANOVA with Šídák’s test, corrected for multiple comparisons. Microscopy images were processed in ImageJ. Background was subtracted, and images were adjusted for brightness and contrast using identical parameters for all images. Quantification of infected macrophages showing positive restaining (HLA-DR, L243) was performed on 20× merged images. The number of HLA-DR (L243) positive infected macrophages (GFP positive) was divided by total number of infected macrophages in 4-5 fields of view and multiplied by 100 to achieve the percentage of infected macrophages with HLA-DR (L243) staining. Percentages of infected macrophages with surface MHC I/II re-staining were compared between conditions using either Student’s t-test or one-way ANOVA with Šídák’s post test, corrected for multiple comparisons respectively. Two-tailed p values < 0.05 were considered significant. For CD4+ T cells, co-expression of CD69 with CD40L or IFN-γ was determined after gating on Live/Dead− CD4+ CD45RA^Lo^ single cells. Statistical analyses and graphs were generated using Prism v10 (GraphPad, San Diego, CA). Data were tested for normality. If a lognormal distribution was detected, a paired mixed-effects one-way ANOVA was used with Geisser-Greenhouse correction and Šídák’s post test, corrected for multiple comparisons. Representative data from individual macrophage experiments were analyzed by ordinary one-way ANOVA with Šídák’s multiple comparisons test. Two-tailed p values < 0.05 were considered statistically significant.

## Data Availability

No new code was generated in this study. Bulk RNA sequencing data used in Figs. 2 and 3 will be publicly available upon publication through the NCBI GEO accession number GSE336443. Comparisons used to generate differentially expressed genes and the gene lists indicated in Figs. 2 and 3 are available as Extended data 1 and Extended data 2, published with the article. Other data will be deposited into the Dryad data repository upon publication or provided by the corresponding author upon reasonable request.

## Acknowledgements

The following reagents were obtained through BEI Resources, NIAID, NIH: *Mycobacterium tuberculosis*, Strain H37Rv, Gamma-Irradiated Whole Cells, NR-49098, and *Mycobacterium tuberculosis* strain H37Rv, NR-13648. Fig. 1A was created in BioRender. This work was supported by National Institutes of Health (NIH) grants K08 AI163407 and R01 AI187662 (to S.M.C.), American Lung Association Innovation Award 1267638 (to S.M.C.), and research funds from University Hospital Cleveland Medical Center (to S.M.C.). Intellectual support was provided by NIH IMPAc-TB Center contract 75N93019C00071. The use of core facilities was supported in part by NIH grant P30 AI036219 to Case Western Reserve University and the University of Pittsburgh Rustbelt Center for AIDS Research (CFAR). Research bronchoscopies were performed by R.F.S. in the Dahms Clinical Research Unit (DCRU), which is a clinical core of the Case Western Reserve University/University Hospitals Cleveland Medical Center Clinical and Translational Science Collaborative, supported by NIH UL1 RR024989 from the National Center for Research Resources. We are grateful to Professors W. Henry Boom and Clifford Harding, and members of the Carpenter Lab for thoughtful scientific feedback.

## Author contributions

Conceptualization, A.K.S., S.M.C.; Data curation, A.K.S., D.P.G., S.M.C.; Formal Analysis, A.K.S., D.P.G., S.M.C.; Funding Acquisition, S.M.C.; Investigation, A.K.S., D.P.G., R.C.S., R.F.S., S.M.C.; Methodology, A.K.S., D.P.G., C.B., B.D.B., R.F.S., S.M.C.; Project Administration, A.K.S., D.P.G., C.B., R.F.S., S.M.C.; Resources, A.K.S., L.H., D.W., C.B., B.D.B., R.F.S., S.M.C.; Software, A.K.S., D.P.G., S.M.C.; Supervision, S.M.C.; Validation, A.K.S., D.P.G., S.M.C.; Visualization, A.K.S., S.M.C.; Writing-original draft preparation, A.K.S., S.M.C.; Writing-review and editing, A.K.S., D.P.G., R.C.S., L.H., D.W., C.B., B.D.B., R.F.S., S.M.C.

## Conflicts of interest statement

The authors have declared that no competing interests exist.

## Supplemental Figures

**Supplemental Figure 1 (related to Figure 3).**
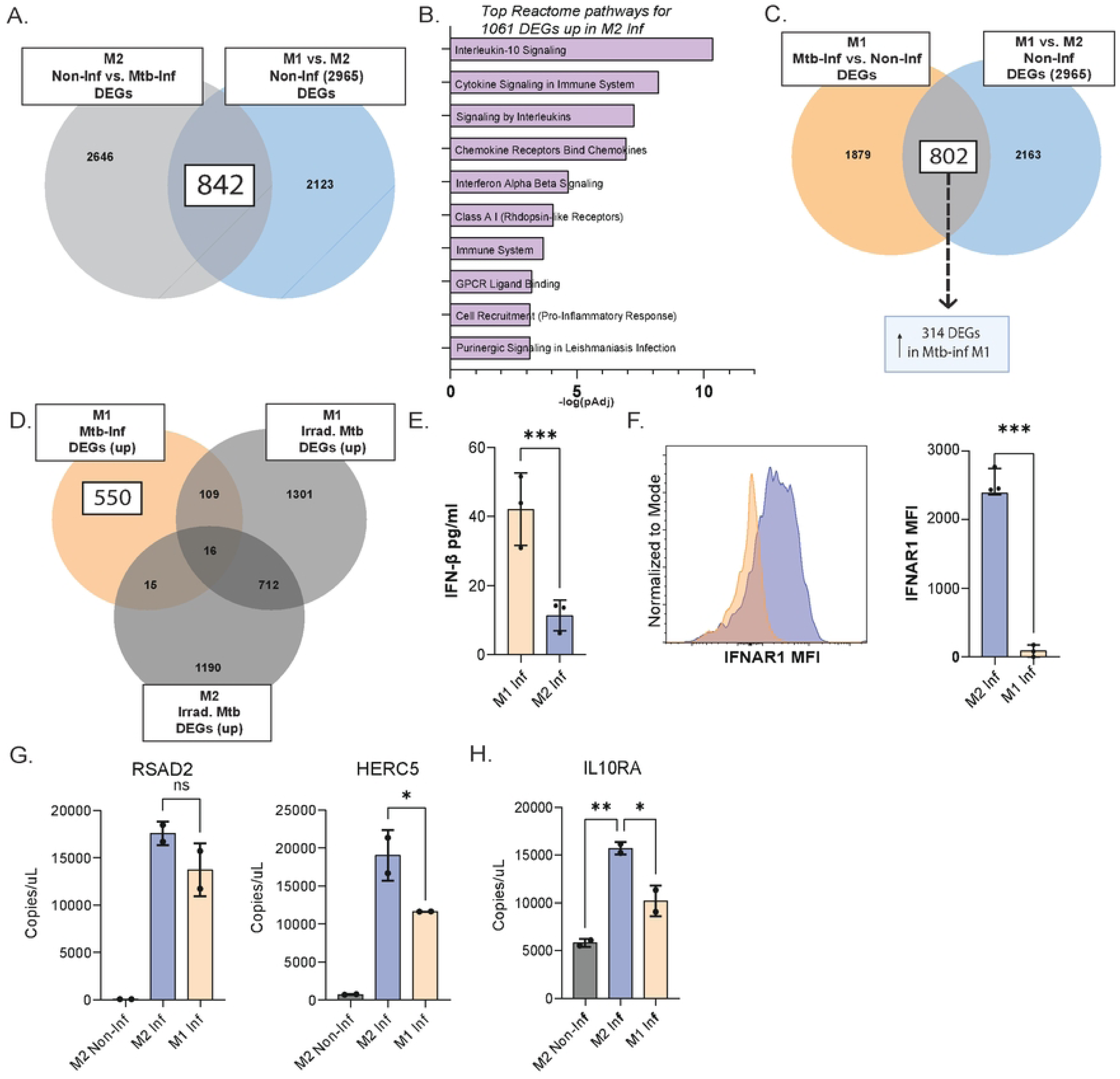
M1-like Mtb-infected macrophages show decreased expression of *HERC5* and *IL10RA.* **(A)** Venn diagram depicting DEGs between non-infected M2-like macrophages and after Mtb infection (light gray) compared to DEGs between non-infected M1 and M2-like macrophages (blue). **(B)** Significant (adjusted) pathways (Reactome) from 1061 genes upregulated by M2-like macrophages after Mtb infection. Pathway enrichment significance was determined by Fisher’s exact test with Benjamini-Hochberg correction for multiple comparisons, as implemented in Enrichr. **(C)** Venn diagrams depicting DEGs between non-infected M1-like macrophages and after Mtb infection (orange) and DEGs between non-infected M1 and M2-like macrophages (blue). **(D)** Venn diagrams depicting DEGs unique to Mtb-infected M1-like macrophages (orange) after removing upregulated DEGs in common with M1 and M2-like macrophages after Irrad. Mtb (compared to non-infected, grey). **(E)** Bar graph showing mean (+/- SD) IFN-β concentrations in supernatants of infected M1 and M2-like macrophages 24 h post infection, in triplicate, representative of 3 independent experiments (3 individuals). **(F) (Left)** Representative histogram and **(right)** bar graph showing MFI (mean + SD) of IFNAR1, in triplicate, in M1 and M2-like MDMs, representative of 2 independent experiments (3 individuals). **(G)** Bar graphs showing the mean mRNA concentrations (+/- SD) of *RSAD2* and *HERC5* and **(H)** *IL10RA* in non-infected and infected M2 and M1-like MDMs, in duplicate, representative of 2 independent experiments (4 individuals). Significance was determined using either Student’s t-test or one-way ANOVA with Sidak’s post-test corrected for multiple comparisons. ∗∗ p < 0.01; ∗∗∗∗p < 0.0001; ns, not significant.

**Supplemental Figure 2 (related to Figure 4).**
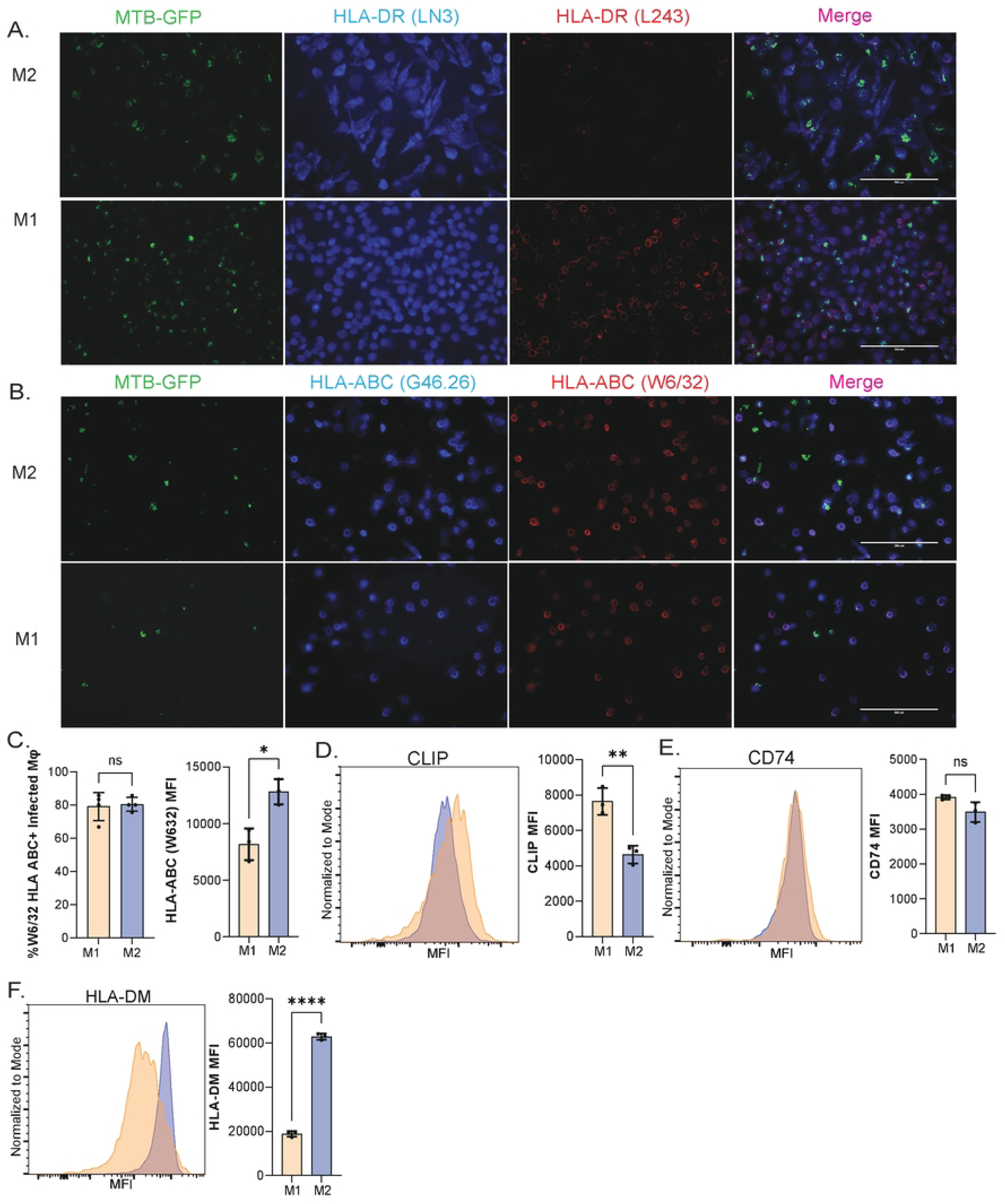
MHC-I trafficking is not restricted in Mtb-infected M2-like macrophages. **(A)** Fluorescence microscopy of M1 and M2-like MDMs treated with anti-HLA-DR mAb blockade (clone L243), infected with Mtb-GFP showing surface anti-HLA-DR staining (BV421-LN3) and re-stained with HLA-DR (PE-L243) 24 h post infection. Images taken at 20x magnification. **(B)** Fluorescence microscopy of M1 and M2-like MDMs treated with anti-HLA-ABC mAb blockade (clone W6/32), infected with Mtb-GFP showing surface anti-HLA-ABC staining (BV421-G46-2.6) and re-stained with HLA-ABC (PE-W6/32) 24 h post infection. Images taken at 20x magnification. **(C) (left)** Bar graph of the percentage (mean ± SD) of infected macrophages showing positive HLA-ABC (PE-W6/32) re-staining, representative of 3 independent experiments (3 individuals) and **(right)** Bar graph showing MFI (mean + SD) of post-infection re-stained HLA-ABC (PE-W6/32) in triplicate, representative of 3 independent experiments (3 individuals). **(D) (left)** Representative histogram and **(right)** bar graph showing MFI (mean + SD) of post-infection HLA-DR-bound surface CLIP in triplicate, representative of 4 independent experiments (2 individuals). **(E) (left)** Representative histogram and **(right)** bar graph showing MFI (mean + SD) of post-infection total CD74 in triplicate, representative of 3 independent experiments (3 individuals). **(F) (left)** Representative histogram and **(right)** bar graph showing MFI (mean + SD) of post-infection total HLA-DM in triplicate, representative of 3 independent experiments (3 individuals). Significance was determined using Student’s t-test. ∗∗ p < 0.01; ∗∗∗∗p < 0.0001; ns, not significant.

**Supplemental Figure 3 (related to Figure 5).**
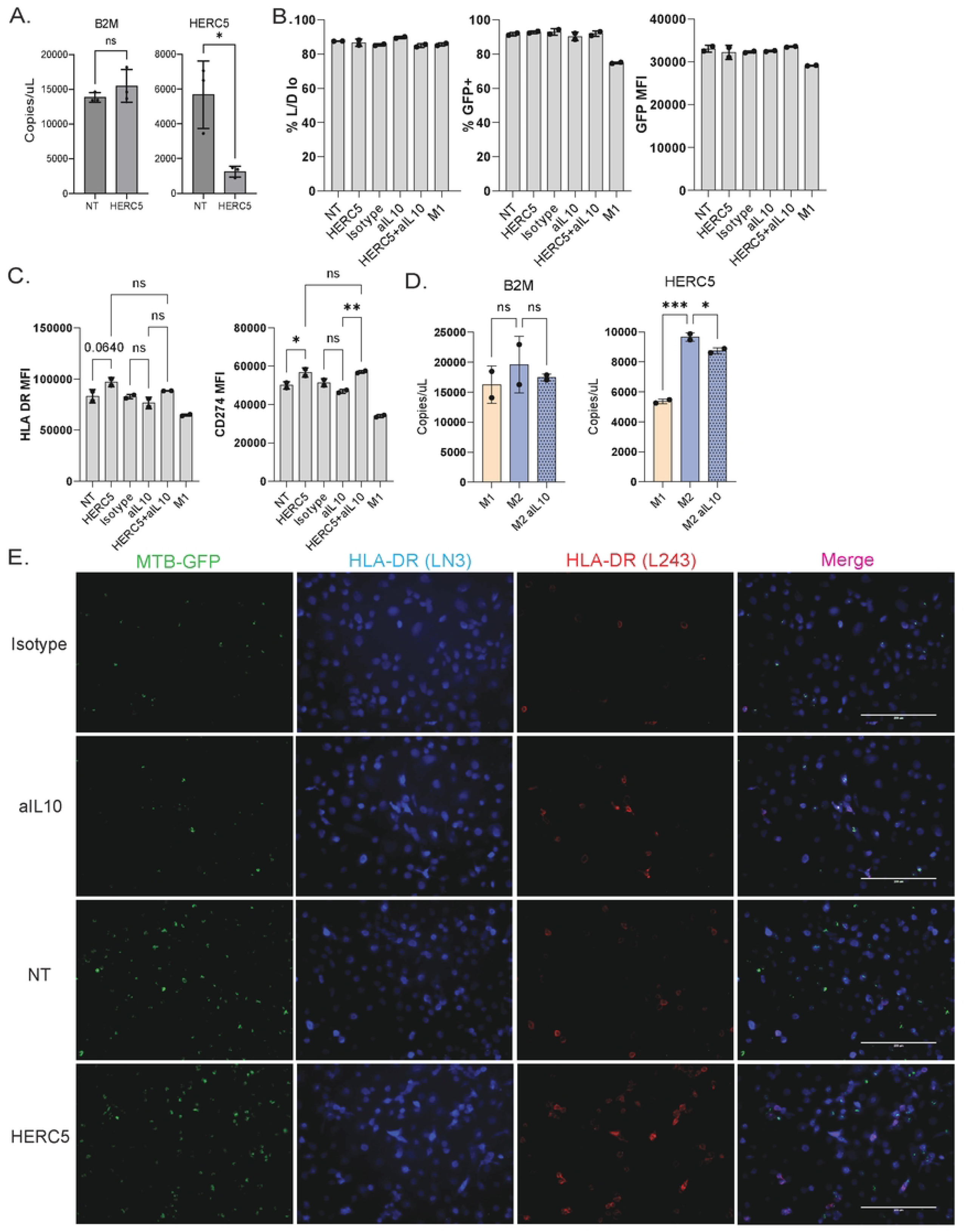
*HERC5* knockdown does not alter infectivity or viability of M2-like macrophages. (**A)** Bar graph showing mean (+ SD) mRNA concentrations of *B2M* **(left)** and *HERC5* **(right)** in infected M2-like macrophages receiving either non-targeting (NT) siRNA or *HERC5* siRNA in triplicate, representative of 3 independent experiments (3 individuals). **(B)** Bar graphs showing mean (+ SD) **(left)** percentage of Live/Dead low M1 and M2-like macrophages under each indicated condition, **(middle)** percentage of Mtb-GFP infected M1 and M2-like macrophages under each indicated condition, and **(right)** MFI of Mtb-GFP in infected M1 and M2-like macrophages under each indicated condition in duplicate, representative of 3 independent experiments (3 individuals). **(C)** Bar graph showing MFI (mean + SD) of post-infection **(left)** surface HLA-DR and **(right)** CD274 in M1 and M2-like macrophages under each indicated condition in duplicate, representative of 3 independent experiments (3 individuals). **(D)** Bar graph showing mean (+ SD) mRNA concentrations of *B2M* **(left)** and *HERC5* **(right)** in infected M1, M2-like macrophages, and infected M2-like macrophages receiving IL-10 neutralization in duplicate, representative of 3 independent experiments (3 individuals). **(E)** Fluorescence microscopy of M2-like MDMs under indicated conditions treated with anti-HLA-DR mAb blockade (clone L243), infected with Mtb-GFP showing surface anti-HLA-DR staining (BV421-LN3) and re-stained with HLA-DR (PE-L243) 24 h post infection. Images taken at 20x magnification. Significance was determined using either Student’s t-test **(A)** or one-way ANOVA with Sidak’s post-test corrected for multiple comparisons **(B, C,** and **D)**. ∗∗ p < 0.01; ∗∗∗∗p < 0.0001; ns, not significant.

**Extended Data 1 (Related to Figures 2 and 3)** includes lists of differentially expressed genes (DEGs) between M1- and M2-like MDMs, and BAL macrophages in pairwise comparisons, as indicated in Figures 2 and 3. **Extended Data 2 (Related to Figures 2 and 3)** includes the gene lists used to construct the Venn diagrams capturing the DEGs in M1- and M2-like MDMs, and BAL macrophages in Figures 2 and 3.

## Notes

### Competing Interest Statement

The authors have declared no competing interest.

